# PMRT1, a *Plasmodium* specific parasite plasma membrane transporter is essential for asexual and sexual blood stage development

**DOI:** 10.1101/2021.12.21.473770

**Authors:** Jan Stephan Wichers, Paolo Mesén-Ramírez, Gwendolin Fuchs, Jing Yu-Strzelczyk, Jan Stäcker, Heidrun von Thien, Arne Alder, Isabelle Henshall, Benjamin Liffner, Georg Nagel, Christian Löw, Danny Wilson, Tobias Spielmann, Shiqiang Gao, Tim-Wolf Gilberger, Anna Bachmann, Jan Strauss

**Affiliations:** Centre for Structural Systems Biology, 22607 Hamburg, Germany; Bernhard Nocht Institute for Tropical Medicine, 20359 Hamburg, Germany; Biology Department, University of Hamburg, 20146, Hamburg, Germany; Institute of Physiology, Department of Neurophysiology, Biocenter, University of Wuerzburg, 97070 Würzburg, Germany; Research Centre for Infectious Diseases, School of Biological Sciences, University of Adelaide, Adelaide 5005, Australia; European Molecular Biology Laboratory, Hamburg Unit, Hamburg, Germany; Burnet Institute, 85 Commercial Road, Melbourne 3004, Victoria, Australia

## Abstract

Membrane transport proteins perform crucial roles in cell physiology. The obligate intracellular parasite *Plasmodium falciparum*, an agent of human malaria, relies on membrane transport proteins for the uptake of nutrients from the host, disposal of metabolic waste, exchange of metabolites between organelles and generation and maintenance of transmembrane electrochemical gradients for its growth and replication within human erythrocytes. Despite their importance for *Plasmodium* cellular physiology, the functional roles of a number of membrane transport proteins remain unclear, which is particularly true for orphan membrane transporters that have no or limited sequence homology to transporter proteins in other evolutionary lineages. Therefore, in the current study, we applied endogenous tagging, targeted gene disruption, conditional knockdown and knockout approaches to investigate the subcellular localization and essentiality of six membrane transporters during intraerythrocytic development of *P. falciparum* parasites. They are localized at different subcellular structures – the food vacuole, the apicoplast, and the parasite plasma membrane – and four out of the six membrane transporters are essential during asexual development. Additionally, the plasma membrane resident transporter 1 (PMRT1, PF3D7_1135300), a unique *Plasmodium*-specific plasma membrane transporter, was shown to be essential for gametocytogenesis and functionally conserved within the genus *Plasmodium*. Overall, we reveal the importance of four orphan transporters to blood stage *P. falciparum* development, which have diverse intracellular localizations and putative functions.

**Importance:** *Plasmodium falciparum*-infected erythrocytes possess multiple compartments with designated membranes. Transporter proteins embedded in these membranes do not only facilitate movement of nutrients, metabolites and other molecules between these compartments, but are common therapeutic targets and can also confer antimalarial drug resistance. Orphan membrane transporter in *P. falciparum* without sequence homology to transporters in other evolutionary lineages and divergent to host transporters may constitute attractive targets for novel intervention approaches. Here, we localized six of these putative transporters at different subcellular compartments and probed into their importance during asexual parasite growth using reverse genetic approaches. In total, only two candidates turned out to be dispensable for the parasite, highlighting four candidates as putative targets for therapeutic interventions. This study reveals the importance of several orphan transporters to blood stage *P. falciparum* development.

## Introduction

*Plasmodium* spp. malaria parasites inhabit diverse intracellular niches and need to import nutrients and export waste across both, host-cell and parasite membranes. Despite this, there are less than 150 putative membrane transporters encoded in the genome of *Plasmodium falciparum*, the most virulent malaria parasite, making up only 2.5% of all encoded genes (*P. falciparum* 3D7 v3.2: 5280 genes) (1–8), which is reduced compared to other unicellular organisms of similar genome size. The loss of redundant transporters is a typical feature of many intracellular parasites (9) and, as a result, the proportion of transporters that are indispensable for parasite survival increases (2), some of which have been shown to be critical for the uptake of several anti-Plasmodial compounds and/or to be involved in drug resistance (10–23). Moreover, the parasite’s intracellular lifestyle resulted in the evolution of additional specialized transporters without human homologues (1). During its intraerythrocytic development, the parasite relies on the uptake of nutrients, such as amino acids, pantothenate or fatty acids, from its host erythrocyte as well as from the extracellular blood plasma (24–27). As *P. falciparum* resides in a parasitophorous vacuole (PV) in the host erythrocyte, nutrients acquired from the extracellular milieu must traverse multiple membranes: the erythrocyte plasma membrane (EPM), the parasitophorous vacuole membrane (PVM), the parasite plasma membrane (PPM) and eventually membranes of intracellular organelles, such as those of the apicoplast or mitochondria (24, 28–30). The unique requirements of malaria parasite survival have led to the evolution of a number of orphan transporters, whose localization or function cannot be predicted based on sequence homology to transporters in other organisms (4, 31). Despite the likely importance of uniquely adapted transporters to *P. falciparum* survival, subcellular localization, essentiality, function and substrate specificity for most *P. falciparum* transporters has not been directly determined (2, 24, 29). The best functional evidence available for many *Plasmodium*-specific transporters comes from a recent knockout screen of these orphan transporters in the rodent malaria parasite *Plasmodium berghei* (31). However, whether observations for different transporters in the *P. berghei* model are directly transferrable to *P. falciparum* have yet to be examined. Therefore, in this study, we explored the localization and essentiality of four predicted orphan transporters that had been partially characterised in *P. berghei* and included two additional transporters with no experimental characterization available.

## Results

To date, the predicted ‘transportome’ of *P. falciparum* consists of 117 putative transport systems (encoded by 144 genes) classified as channels (n=19), carriers (n=69), and pumps (n=29) (2). Functions of the vast majority of transporter genes were inferred from sequence homology to model organisms, however, given their lack of homology, 39 gene products could not be associated with any functional or subcellular localization and were categorized as orphan transporters accordingly (4). A subset of orphan transporters characterized in the *P. berghei* malaria model was selected for further characterization in *P. falciparum*. The four transporters selected were reported to be important at different stages of rodent malaria parasite growth with i) *P. berghei* drug/metabolite transporter 2 (*Pf*DMT2: PF3D7_0716900) found to be essential for asexual blood stage development, ii) *P. berghei* zinc transporter 1 (*Pf*ZIP1: PF3D7_0609100) was essential across transmission stages but not blood stages, where there was only a slight growth defect, iii) *P. berghei* cation diffusion facilitator family protein (*Pf*CDF: PF3D7_0715900) knockout parasites had a defect during transmission stages but not during asexual stages, and iv) *P. berghei* major facilitator superfamily domain-containing protein (*Pf*MFS6: PF3D7_1440800) was found to be essential for parasite transmission from mosquitos to a new host, with a growth defect observed at asexual and gametocyte stages but not during mosquito stage parasite growth (31, 32). In order to confirm expression of these four, initially selected, transporters in *P. falciparum* asexual stages, we searched the list of “Genes coding for transport proteins” included in the Malaria Parasite Metabolic Pathways (MPMP) database (1, 33) for proteins with i) RNA-seq (34, 35) and ii) proteomics evidence (36, 37) in asexual blood stages. During our initial searches of the MPMP database but also including PlasmoDB (38) and the most recent *P. falciparum* 3D7 genome (v3.2) and annotations, we identified two additional putative transporters in *P. falciparum* (PF3D7_0523800, PF3D7_1135300), whose *P. berghei* homologs were not targeted and functionally characterized by Kenthirapalan *et al*. (31) or investigated in any other experimental model. Given their obvious lack of sequence homology to transporter proteins in other evolutionary lineages and clear classification as orphan membrane transporter, both proteins were subsequently included in our characterization of *P. falciparum* orphan transporters, and named as ‘food vacuole resident transporter 1’ (FVRT1: PF3D7_0523800) and as ‘plasma membrane resident transporter 1’ (PMRT1: PF3D7_1135300) based on their subcellular localization. AlphaFold-based structure predictions (39) and results from structure homology search (40) of all six selected transporters are provided in Figure S1.

### Localization of putative *P. falciparum* transporters

To determine subcellular localization, we tagged the six putative transporters endogenously with GFP using the selection-linked integration (SLI) system (41) (Figure 1A). Additionally, a glmS ribozyme sequence was included in the 3’UTR, which enabled conditional gene knockdown upon addition of glucosamine (42). Correct integration of the plasmid into the respective genomic locus was verified by PCR and expression of the GFP-fusion protein was confirmed by Western blot for each generated cell line (Figure S2A, B).

**Figure 1:**
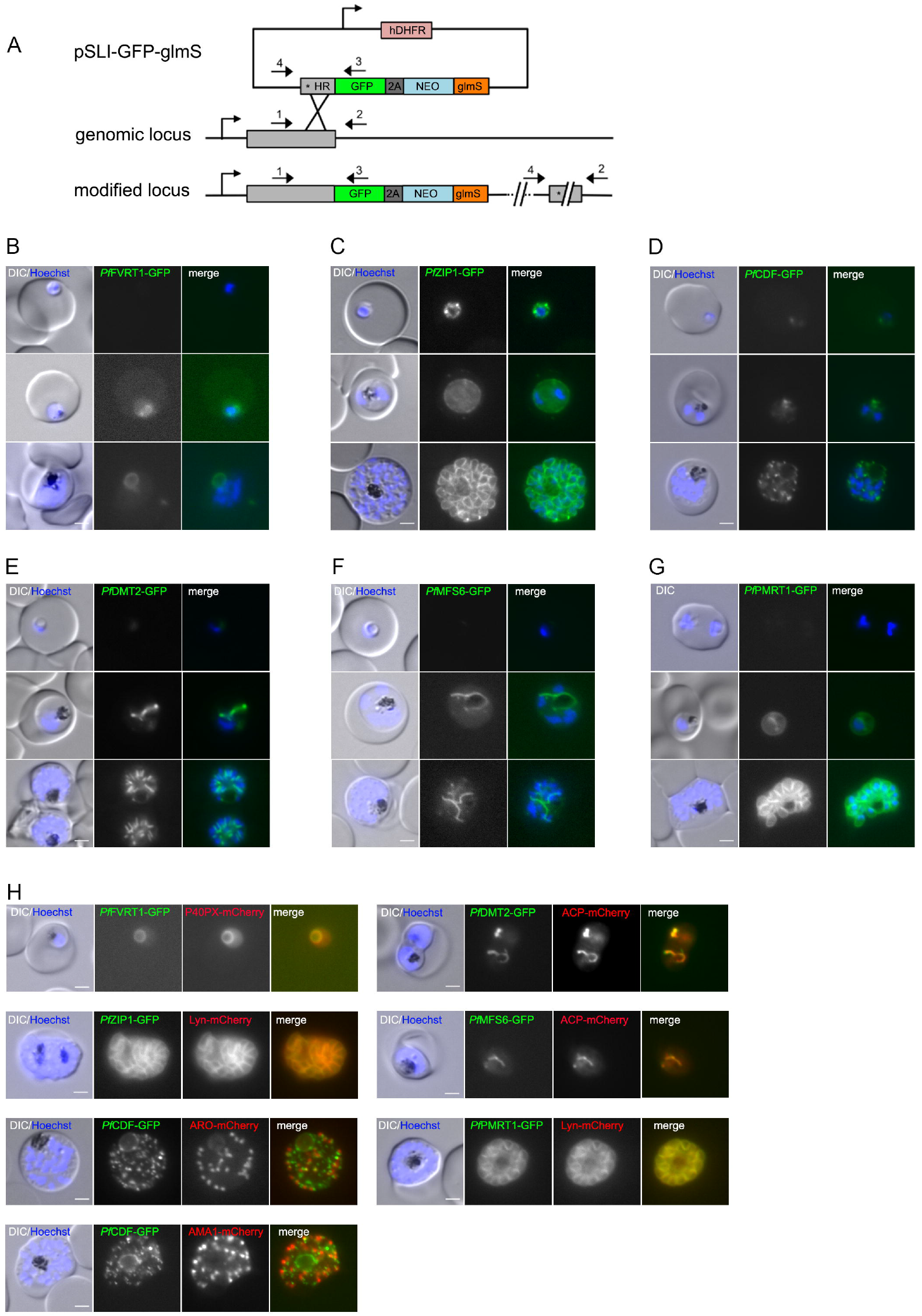
Subcellular localization of six putative *P. falciparum* transporters during asexual blood stage development. **(A)** Schematic representation of endogenous tagging strategy using the selection-linked integration system (SLI). pink, human dihydrofolate dehydrogenase (hDHFR); grey, homology region (HR); green, green fluorescence protein (GFP) tag; dark grey, T2A skip peptide; blue, neomycin resistance cassette; orange, glmS cassette. Stars indicate stop codons, and arrows depict primers (P1 to P4) used for the integration check PCR. **(B–G)** Localization of (B) *Pf*FVRT1-GFP-glmS, (C) *Pf*ZIP1-GFP-glmS, (D) *Pf*CDF-GFP-glmS, (E) *Pf*DMT2-GFP-glmS, (F) *Pf*MFS6-GFP-glmS and (G) *Pf*PMRT1-GFP-glmS by live-cell microscopy in ring, trophozoite and schizont stage parasites. Nuclei were stained with Hoechst-33342. **(H)** Co-localization of the GFP-tagged putative transporters with marker proteins P40PX-mCherry (food vacuole), ACP-mCherry (apicoplast), Lyn-mCherry (parasite plasma membrane), ARO-mCherry (rhoptry) and AMA1-mCherry (microneme) as indicated. Nuclei were stained with Hoechst-33342. Scale bar, 2 µm.

All transgenic cell lines expressed the GFP-fusion protein, demonstrating that these transporters are expressed in asexual blood stage parasites (Figure 1B-G, S2A). Expression levels were sufficient to allow determination of subcellular localization (Figure 1B–G): (i) PF3D7_0523800-GFP localized to the food vacuole, (ii) *Pf*DMT2-GFP and *Pf*MFS6-GFP apicoplast localization, and (iii) *Pf*ZIP1-GFP and PF3D7_1135300-GFP parasite plasma membrane (PPM) localization. However, *Pf*CDF-GFP showed an obscure staining pattern with a weak spot within the parasite cytosol in ring and trophozoite state parasites, but multiple foci in schizont stages (Figure 1D). To pinpoint this localization, an additional cell line with endogenously 3xHA-tagged *Pf*CDF was generated, confirming the focal localization of *Pf*CDF in asexual stages (Figure S2C).

Except for *Pf*CDF, the observed localizations of the other five transporters were confirmed by co-localization studies using appropriate episomally expressed marker proteins: P40PX-mCherry (43, 44) for the food vacuole, ACP-mCherry (45, 46) for apicoplast and Lyn-mCherry (41, 47) for PPM. The focal distribution of *Pf*CDF-GFP was co-localized with a rhoptry (ARO-mCherry (48, 49)) and a micronemes (AMA1-mCherry (50, 51)) marker, but *Pf*CDF-GFP did not colocalize with either marker (Figure 1H). Additionally, for *Pf*ZIP and PF3D7_1135300 the PPM localization was further confirmed in free merozoites (Figure S2D, E) and by confocal microscopy-based co-localization of PF3D7_1135300-GFP with the PPM marker Lyn-mCherry (Figure S2F). Accordingly, as noted above, we named PF3D7_0523800 as ‘food vacuole resident transporter 1’ (FVRT1) and PF3D7_1135300 as ‘plasma membrane resident transporter 1’ (PMRT1).

### Targeted-gene disruption (TGD), conditional knockdown and conditional knockout of putative transporters

In order to test whether the putative transporters are essential for *P. falciparum* during its intraerythrocytic cycle, we first tried to functionally inactivate them by targeted gene disruption (TGD) using the SLI system (41) (Figure S3A). TGD cell lines were successfully obtained for *Pf*ZIP1 and *Pf*CDF (Figure S3B, C). For *Pf*ZIP1-TGD, the correct integration of the plasmid into the genomic locus and absence of wildtype locus was verified by PCR and subsequent growth experiments revealed no growth defect compared to *P. falciparum* 3D7 wildtype parasites (Figure S2B), suggesting its redundancy during asexual parasite proliferation. For *Pf*CDF-TGD the correct integration of the plasmid into the genomic locus was also verified, but wildtype DNA was still detectable and remained even upon prolonged culturing under G418/WR selection and limited dilution cloning (Figure S3C). In contrast, six (*Pf*PMRT1, *Pf*DMT2) or eight (*Pf*FVRT1, *Pf*MFS6) independent attempts to obtain TGD cell lines for the other four transporters with the respective plasmids failed, indicating that these genes have an indispensable role in blood stage parasite growth.

To probe into the function of the putative transporters where we were unable to generate gene-disruptions, we utilized the glmS ribozyme sequence. The corresponding sequence was integrated into the 3’UTR of the targeted genes. This enabled the induction of conditional degradation of respective mRNAs upon addition of glucosamine (42) and the assessment of the phenotypic consequences. Upon addition of 2.5 mM glucosamine to young ring stage parasites we found a 76.8% (+/- SD 3.7) reduction in GFP fluorescence intensity in *Pf*DMT2GFP parasites, 72.7% (+/- SD 9.4) reduction in *Pf*MFS6-GFP and a 77.7% (+/- SD 6.1) reduction in *Pf*PMRT1-GFP in schizonts of the same cycle (Figure 2A–C, S4A–C). No measurable reduction in fluorescence intensity could be detected for *Pf*FVRT1-GFP or *Pf*CDF-GFP expressing parasite lines (Figure S4D–F). Presence of the glmS cassette in both plasmids was confirmed by PCR (Figure S4H). For parasite cell lines with a significant reduction in the expression of the endogenously tagged protein, proliferation was analyzed in the absence and presence of 2.5 mM glucosamine (Figure 2D, S4G). While no significant effect on growth was observed for *Pf*MFS6, a growth reduction of 68.5 % (+/- SD 2.1) over two cycles was observed upon knockdown of *Pf*DMT2. For *Pf*PMRT1, a minor growth delay was measurable, which resulted in a significantly reduced parasitemia at day 3 upon knockdown using 2.5 mM glucosamine (two tailed Wilcoxon rank sum test, *W* = 15, *n*_1_ = 5, *n*_2_ = 3, *P* = 0.03), but was not significant when using 5 mM glucosamine (two tailed Wilcoxon rank sum test, *W* = 10, *n*_1_ = 4, *n*_2_ = 3, *P* = 0.16) (Figure 2E). Additionally, significantly fewer newly formed ring stage parasites were observed at 84 hours post invasion (hpi) (Figure 2F), and multiple pairwise post-hoc comparisons using the Conover-Iman rank sum test and Benjamini-Hochberg method to control the false discovery rates showed significant step-wise reductions of ring stage parasites after induction of GlmS-based knockdown of PfPMRT1 using both, 2.5 mM glucosamine (adjusted *P* = 0.0078) and 5 mM glucosamine (adjusted *P* = 0.0005) in comparison to untreated control cell cultures.

**Figure 2:**
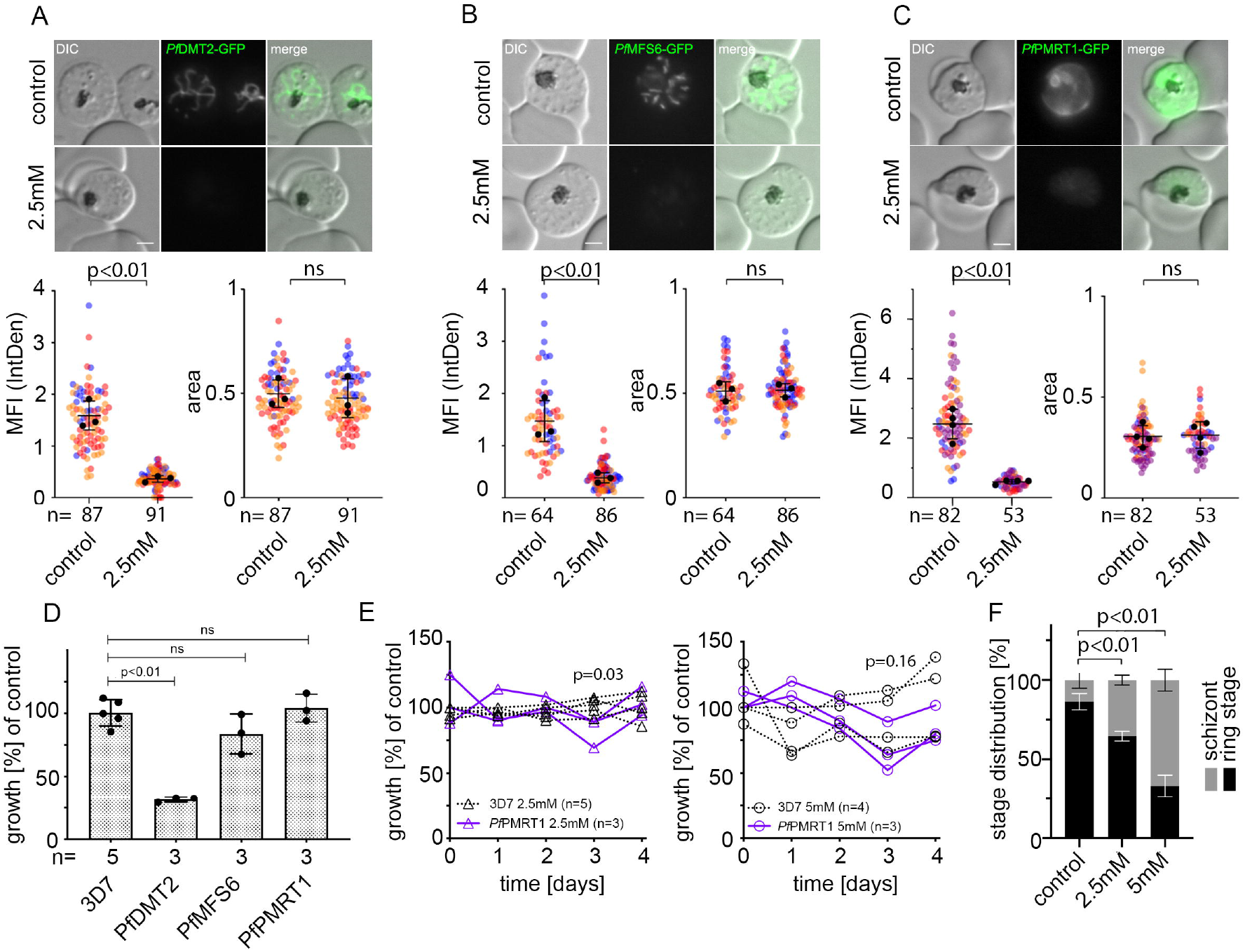
Conditional knockdown of putative transporter indicate importance of *Pf*DMT2 and *Pf*PMRT1 for parasites fitness. **(A–C)** Live cell microscopy and quantification of knockdown by measuring mean fluorescence intensity (MFI) density and size (area) of (A) *Pf*DMT2-GFP-glmS (B) *Pf*MFS6-GFP-glmS and (C) *Pf*PMRT1-GFP-glmS parasites 40 hours after treatment without (control) or with 2.5 mM glucosamine. Scale bar, 2 µm. Statistics are displayed as mean +/- SD of three (A–B) or four (C) independent experiments and individual data points are color-coded by experiments according to Superplots guidelines (101). P-values displayed were determined with two-tailed unpaired t-test. **(D)** Growth of parasites treated without (control) or with 2.5 mM glucosamine determined by flow cytometry is shown as relative parasitemia values after two cycles. Shown are means +/- SD of three (*Pf*PMRT1-GFP-glmS, *Pf*DMT2-GFP-glmS, *Pf*MFS6-GFP-glmS) and five (3D7 wild type parasites) independent growth experiments. P-values displayed were determined with unpaired t test with Welch correction and Benjamin-Hochberg for multiple testing correction. Individual growth curves are shown in Figure S4G. **(E)** Growth of *Pf*PMRT1-glmS and 3D7 parasites after treatment with 2.5 mM (left panel) and 5 mM glucosamine (right panel) compared to untreated control parasites over five consecutive days. *P*-values displayed were determined for comparison between *Pf*PMRT1-glmS and 3D7 parasites at day 3 using two-tailed Wilcoxon rank sum test. **(F)** Mean +/- SD distribution of ring and schizont stage parasites in *Pf*PMRT1-glmS and 3D7 cell lines treated without (control), with 2.5 mM or 5 mM glucosamine at 84 hpi (80 hours post addition of glucosa-mine) of three independent experiments. *P*-values displayed were determined using the Conover-Iman rank sum test and Benjamini-Hochberg method for multiple testing correction after Kruskal-Wallis testing.

To better characterize the minor growth phenotype of *Pf*PMRT1-GFP-glmS parasites that might be due to incomplete knockdown, we generated a conditional *Pf*PMRT1 knockout cell line (condΔPMRT1) using the Dimerizable Cre (DiCre) system (52, 53). Again using the SLI system (41), the endogenous *Pf*PMRT1 was disrupted upstream of the region encoding the N-terminal transmembrane domain, but, at the same time introducing a recodonized second functional copy of *Pf*PMRT1 flanked by loxP sites in the genomic locus. This loxP-flanked allelic copy of *Pf*PMRT1 encodes an additional 3x hemagglutinin (HA) tag, which can be conditionally excised upon addition of a rapamycin analog (rapalog) via the enzymatic activity of an episomally expressed DiCre (Figure 3A). First, correct integration of the plasmid into the genomic locus was verified by PCR (Figure 3B). Second, expression and localization of the recodonized HA-tagged protein at the PPM was verified by colocalization with the merozoite plasma membrane marker MSP1 (54) (Figure 3C). Third, excision of the recodonized gene upon rapalog addition was confirmed on genomic level by PCR (Figure 3D) and on protein level by Western blot analysis at 24 hpi and 48 hpi (Figure 3E). To assess the effect of conditional *Pf*PMRT1 knockout on parasite proliferation, we determined growth of the transgenic parasite cell line with and without rapalog over five days (Figure 3F, S5A). In contrast to the glmS-based knockdown experiment, DiCre-based gene excision (induced by the addition of rapalog to young ring stages of condΔPMRT1 parasite cell cultures) abolished growth within the first replication cycle (Figure 3F, S5A). The specificity of the observed growth phenotype was verified by gene complementation. To achieve this, we episomally expressed recodonized *Pf*PMRT1 with TY1-epitope tag either under the constitutive *nmd3* or the weaker *sf3a2* promoter (55) in the condΔPMRT1 cell line (Figure 3D, F, S5B, C). Correct localization of the TY1-tagged *Pf*PMRT1 at the PPM was verified by immunofluorescence assays (IFA) (Figure 3G). Notably both, complementation of the *Pf*PMRT1 knockout cell line (condΔPMRT1) with recodonized *Pf*PMRT1 either under control of the constitutive *nmd3* or the weaker *sf3a2* promoter, restored parasite growth (Figure 3F, S5B, C). The level of growth restoration with low level expression of recodonized *Pf*PMRT1 is in line with the results from glmS-knockdown experiments, which showed that a reduction of about 75% in protein expression resulted only in a minor growth perturbation (Figure 2C, D).

**Figure 3:**
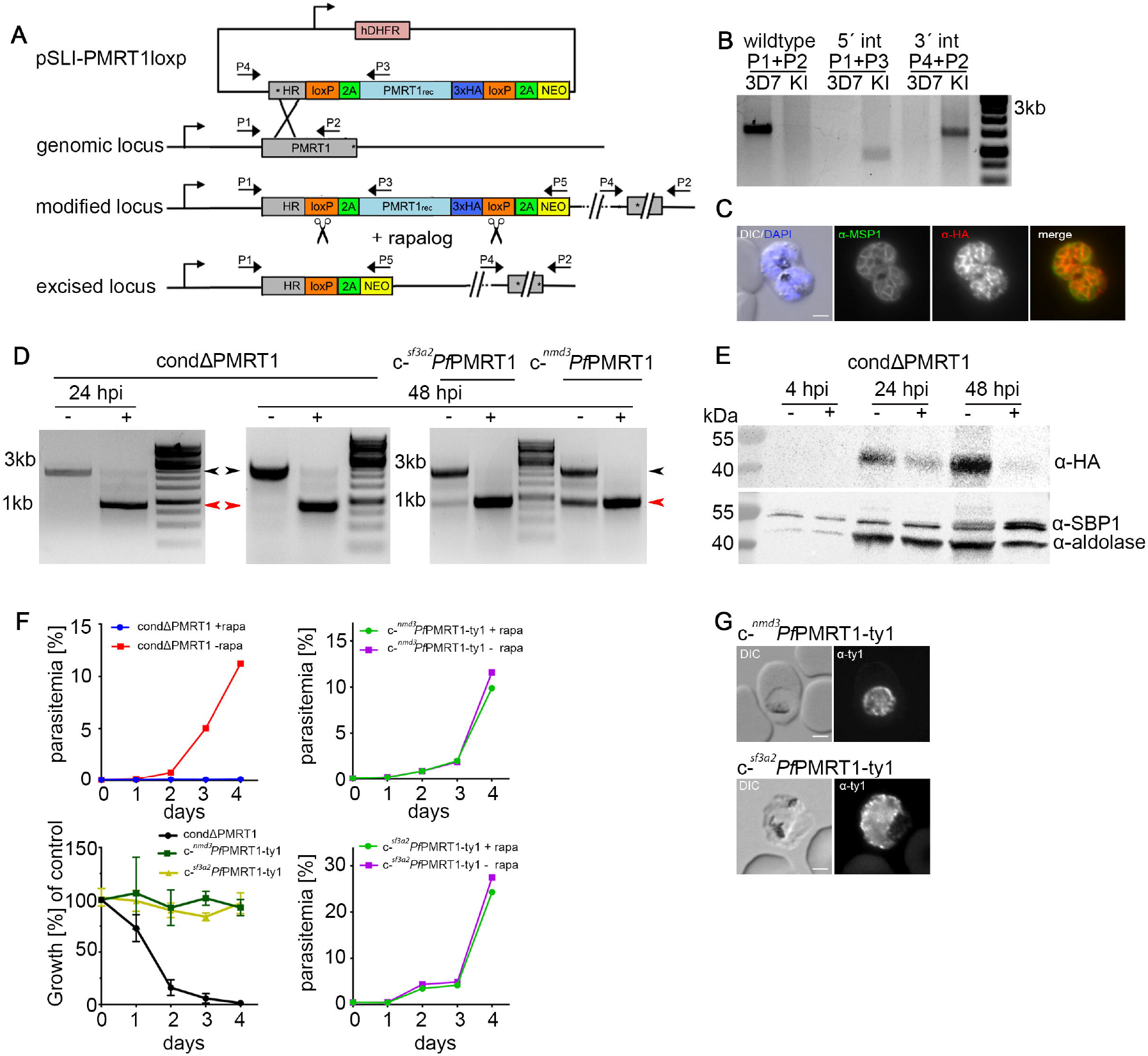
*Pf*PMRT1 is essential for asexual blood stage development. **(A)** Simplified schematic of DiCre-based conditional *Pf*PMRT1 knockout using selection-linked integration (SLI). Pink, human dihydrofolate dehydrogenase (hDHFR); grey, homology region (HR); green, T2A skip peptide; light blue, recodonized *Pf*PMRT1; dark blue, 3xHA tag, yellow, neomycin phosphotransferase resistance cassette; orange, loxp sequence. Scissors indicate DiCre mediated excision sites upon addition of rapalog. Stars indicate stop codons, and arrows depict primers (P1 to P5) used for the integration check PCR and excision PCR. **(B)** Diagnostic PCR of unmodified wildtype and transgenic condΔPMRT1 knock-in (KI) cell line to check for genomic integration using Primer P1-P4 as indicated in (A). **(C)** Immunofluo-rescence assay (IFA) of condΔPMRT1 late stage schizont parasites showing localization of *Pf*PMRT1-3xHA at the parasite plasma membrane (PPM) co-localizing with the merozoite surface protein 1 (MSP1). **(D)** Diagnostic PCR to verify the excision at genomic level at 24 hpi / 20 hours post rapalog addition for condΔPMRT1 and at 48 hpi for condΔPMRT1, c-^*nmd3*^*Pf*PMRT1-ty1 and c-^*sf3a2*^*Pf*PMRT1-ty1 parasites using Primer P1-P5 as indicated in (A). Black arrow head, original locus; red arrow head, excised locus. **(E)** Western blot using α-HA to verify knockout of *Pf*PMRT1 on protein level 4, 24 and 48 hours post invasion. Expected molecular weight of *Pf*PMRT1-3xHA: 53.3 kDa. Antibodies detecting Aldolase and SBP1 were used as loading controls. **(F)** Growth curves of condΔPMRT1, c-^*nmd3*^*Pf*PMRT1-ty1 and c-^*sf3a2*^*Pf*PMRT1-ty1 parasites +/- rapalog monitored over five days by flow cytometry. One representative growth curve is depicted (replicates in Figure S5). Summary is shown as relative parasitemia values, which were obtained by dividing the parasitemia of rapalog treated cultures by the parasitemia of the corresponding untreated ones. Shown are means +/- SD of three (condΔPMRT1, c-^*nmd3*^*Pf*PMRT1-ty1) or four (c-^*sf3a2*^*Pf*PMRT1-ty1) independent growth experiments. **(G)** IFA of condΔPMRT1 complemented with C-terminal TY1-tagged *Pf*PMRT1 constructs expressed either under the constitutive *nmd3* or the weak *sf3a2* promoter to verify PPM localization. Scale bar, 2 µm.

### Loss of the PPM-localized *Pf*PMRT1 leads to an arrest of parasite development at trophozoite stage and the formation of PPM derived protrusions

To determine, which particular parasite stages are affected by the knockout of *Pf*PMRT1, we added rapalog to tightly synchronized parasites at different time points (4, 20 and 32 hpi) (Figure 4A) and monitored parasite growth by flow cytometry. Additionally, we quantified growth perturbation by microscopy of Giemsa smears at 4, 20, 24, 32, 40, 48, 72 and 96 hpi (Figure 4B, S6A, B). When adding rapalog at 4 hpi, parasite development progressed through ring and early trophozoite stages up to 24 hpi with no visible abnormality. Afterwards, parasites with deformed and enlarged protrusions started to appear and further development occurred to be stalled. At 32 hpi, almost all parasites had developed to late trophozoites/early schizonts in the control, whereas these stages were completely absent in *Pf*PMRT1-deficient parasites. Over 50% of the parasites were pycnotic or possessed large protrusions, the remaining parasites stayed arrested at the trophozoite stage. Quantification of the percentage of parasites with protrusions between 20 hpi and 32 hpi revealed 94.8% (+/- SD 4.0) protrusion-positive parasites (Figure 4C). The activation of gene excision at later time points by adding rapalog at 20 hpi or 32 hpi resulted in no or minor growth perturbation in the first cycle with successful re-invasion, but again led to parasites arresting at the trophozoite stage in the second cycle with an accumulation of protrusions (Figure 4A, S6A, B).

**Figure 4:**
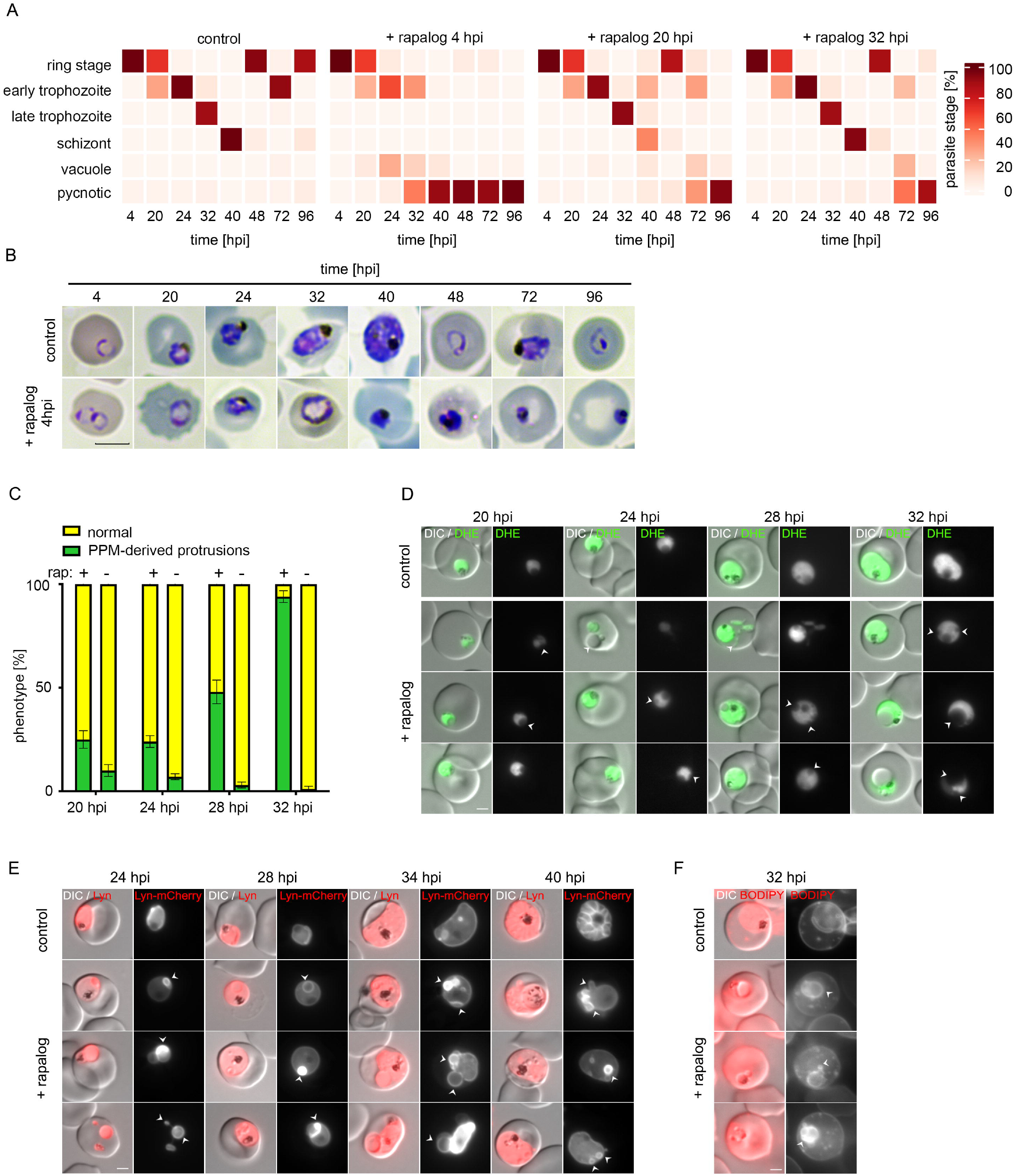
Knockout of *Pf*PMRT1 results in accumulation of PPM-derived protrusions and growth arrest at the trophozoite stage. **(A)** Parasite stage distribution in Giemsa smears displayed as heatmap showing percentage of parasite stages for tightly synchronized (+/- 2 h) 3D7 control and condΔPMRT1 (rapalog treated at 4 hpi, 20 hpi or 32 hpi) parasite cultures over two consecutive cycles. A second replicate is shown in Figure S6A **(B)** Giemsa smears of control and at 4 hpi rapalog treated condΔPMRT1 parasites over two cycles. Scale bar, 5 µm. **(C)** Live cell microscopy of 4 hour window synchronized 3D7 control and condΔPMRT1 parasites +/- rapalog stained with dihydroethidium (DHE) at 20–32 hpi. **(D)** Quantification of parasites displaying protrusions (green) for 4 hour window synchronized 3D7 control and rapalog treated condΔPMRT1 parasites. Shown are percentages of normal parasites versus parasites displaying protrusions as means +/- SD of three independent experiments. **(E)** Live cell microscopy of 8 hour window synchronized 3D7 control and rapalog treated condΔPMRT1 parasites, episomally expressing the PPM marker Lyn-mCherry at 24–40 hpi. **(F)** Live cell microscopy of 3D7 control and condΔPMRT1 parasites +/- rapalog stained with BODIPY TR C5 ceramide at 32 hpi. Scale bar, 2 µm.

In order to get further insights into the morphological changes in *Pf*PMRT1-deficient parasites, we incubated these parasites with dihydroethidium (DHE) to visualize the parasite cytosol (44). We observed an absence of staining within the protrusions, suggesting they are not filled with parasite cytosol (Figure 4D). Next, we transfected the condΔPMRT1 cell line with a plasmid encoding the PPM marker Lyn-mCherry (41) and observed Lyn-mCherry-positive protrusions upon knockout of *Pf*PMRT1 starting to become visible at 24 hpi, indicating that the protrusions originate from the PPM (Figure 4E). In line with this, protrusion membranes were also stainable with BODIPY TR C5 ceramide in condΔPMRT1 parasites at 32 hpi (Figure 4F).

### Depletion of *Pf*PMRT1 results in an early arrest of gametocyte development

RNA-seq data suggest *Pf*PMRT1 is also expressed during other developmental stages, such as gametocytes (56, 57). Therefore, we assessed expression of *Pf*PMRT1-GFP during gametocytogenesis by re-engineering *Pf*PMRT1-GFP-glmS in the inducible gametocyte producer (iGP) ‘3D7-iGP’ (58) parasite line, which allows the robust induction of sexual commitment by conditional expression of gametocyte development 1 protein (GDV1) upon addition of shield-1 (58) (Figure S7A). We show that *Pf*PMRT1 is indeed expressed during all stages of gametocytogenesis and again localizes to the PPM, colocalizing with the PPM-marker Lyn-mCherry (41) (Figure 5A, B). Conditional knockdown of *Pf*PMRT1 via the glmS-ribozyme system (Figure S7B) resulted in a reduction in *Pf*PMRT1-GFP fluorescence intensity of 79.4% (+/- SD 9.2%) at 7 days post induction (dpi) or 75.5% (+/- SD 23.2%) at 10 dpi, without an effect on gametocyte development (Figure S7C–F). In order to exclude that a role of *Pf*PMRT1 in gametocytogenesis is covered by only a partial knockdown resulting in low levels of expressed protein and to determine if *Pf*PMRT1 is essential for gametocytogenesis, we episomally expressed GDV1-GFP-DD in the condΔPMRT1 parasite line, enabling conditional induction of sexual commitment upon addition of shield 1 in these parasites (59). Conditional knockout of *Pf*PMRT1 in these transgenic parasites at day three post gametocyte induction resulted in pycnotic parasites from day 5 onwards, while excision of *Pf*PMRT1 at day 5 post induction had no effect on gametocyte development (Figure 5C, D). Excision of the recodonized gene upon rapalog addition was confirmed at a genomic level by PCR for both conditions (Figure 5E). Quantification of parasite stages at day 10 post induction of GDV1 expression revealed 77.9% (+/- SD 7.7%) gametocytes and 22.1% (+/- SD 7.7%) pycnotic parasites in the control, while 100% of parasites were already pycnotic in the cultures, with induced knockout by addition of rapalog at day 3 post gametocyte induction by GDV1 expression (Figure 5F). This data indicates that *Pf*PMRT1 is important for early gametocyte development.

**Figure 5:**
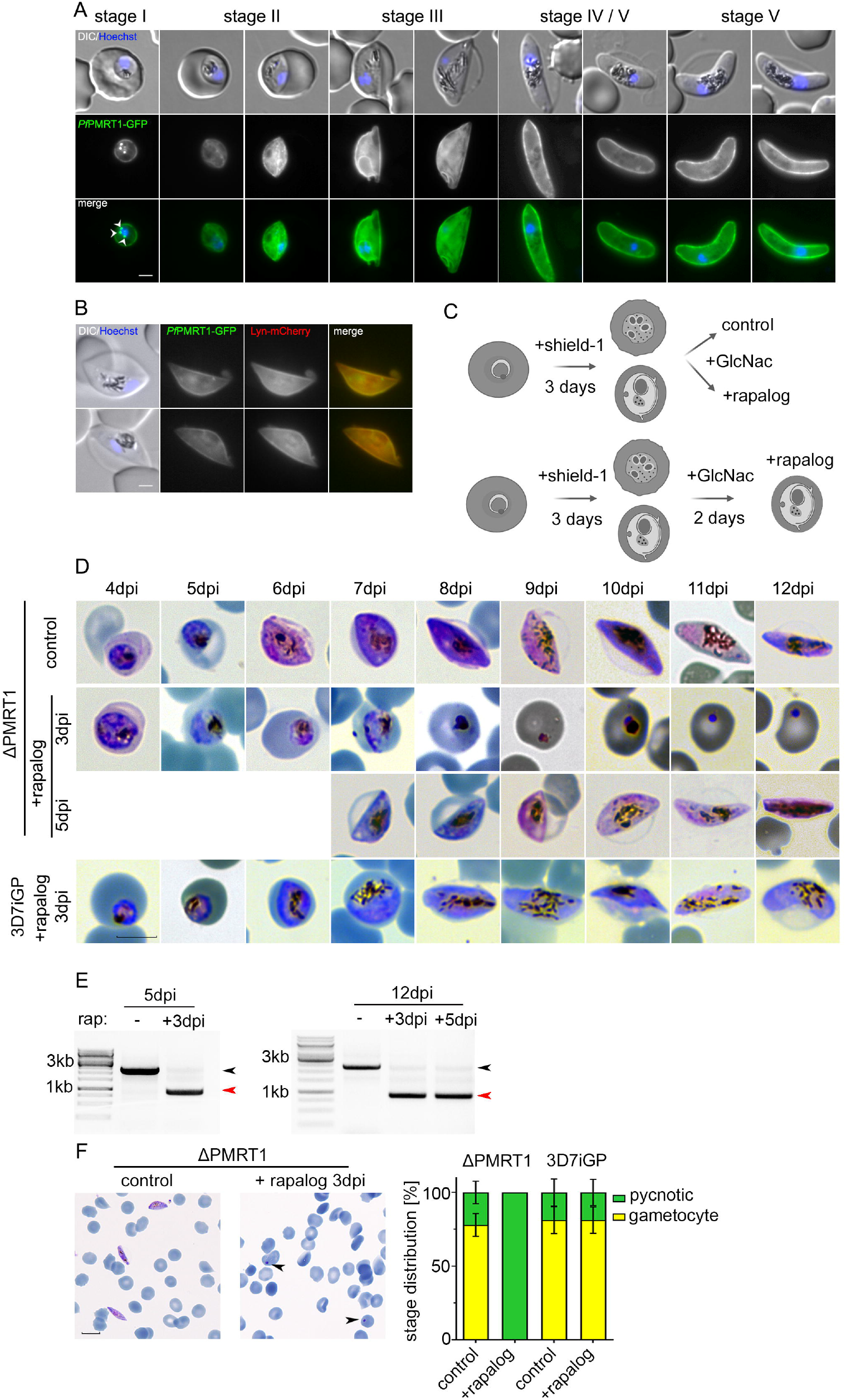
*Pf*PMRT1 is essential for early gametocyte development. **(A)** Live cell microscopy of 3D7-iGP-*Pf*PMRT1-GFP parasites across the complete gametocyte development. White arrow heads indicate remaining GDV1-GFP signal observed in close proximity to the Hoechst signal, as previously reported (59, 94, 102, 103). **(B)** Live cell microscopy of *Pf*PMRT1-GFP parasites expressing the PPM marker Lyn-mCherry. Nuclei were stained with Hoechst-33342. Scale bar, 2 µm. **(C)** Experimental setup of gametocyte induction upon GDV1-GFP-DD expression (+shield-1) and conditional *Pf*PMRT1 knockout (+rapalog) and elimination of asexual blood stage parasites (+GlcNac). **(D)** Gametocyte development over 12 days of condΔPMRT1/GDV1-GFP-DD or 3D7-iGP parasites without (control) or with rapalog addition at day 3 (3 dpi) or day 5 (5 dpi) after induction of sexual commitment by conditional expression of GDV1-GFP upon addition of shield-1. Scale bar, 5 µm. **(E)** Diagnostic PCR to verify the excision on genomic level at 5 dpi and 12 dpi. Black arrow head, original locus; red arrow head, excised locus. **(F)** Representative Giemsa smears and quantification of parasite stage distribution at day 10 post induction for parasites treated without (control) or with rapalog at day 3 post induction. For each condition parasitemia and parasite stages distribution in (ΔPMRT1: n_control_= 3370, 2304, 2759 and n_rapalog_ = 3010, 1830, 2387; 3D7-iGP: n_control_= 4985, 4685, 5206 and n_rapalog_ = 4930, 4332, 5384) erythrocytes of three independent experiments were determined and are displayed as percentage. Nuclei were stained with Hoechst-33342. Scale bar, 10 µm.

### PMRT1 is unique to the genus *Plasmodium* and interspecies complementation assays showed partial functional conservation

*Pf*PMRT1 shows a lack of sequence similarities with known or putative transporters and/or conserved domains shared with known transporter families (2, 5). Our phylogenetic analysis revealed that homologs of *Pf*PMRT1 are present across *Plasmodium* species with amino acid sequence identities of about 90% in the subgenus *Laverania*, but about 50% outside *Laverania* (Figure 6A). However, prediction of the protein structure using AlphaFold (39) indicates two bundles of four transmembrane helices with reasonable similarity of the C-terminal bundle with the photosynthetic reaction center Maquette-3 protein (60) (RMSD of 3.12) (Figure 6B, Figure S1B). In order to test for functional conservation, we expressed the *Pf*PMRT1 homologs of *P. vivax* (PVP01_0936100) and *P. knowlesi* (PKNH_0933400) episomally as C-terminal Ty-1 fusion proteins under the *nmd3* promoter in the condΔPMRT1 parasites. Both fusion proteins are expressed. They were again localized at the PPM as shown by IFA (Figure 6C, Figure S8), and, importantly, were able to partially restore growth after two cycles to 64.8% (+/- SD 9.8%) and 65.1% (+/- SD 7.4%) compared to condΔPMRT1 parasites (Figure 6D, S8). Excision of the recodonized endogenous *Pfpmrt1* gene upon rapalog addition was confirmed at a genomic level by PCR (Figure 6E). These data indicate that PMRT1 is functionally conserved within the genus *Plasmodium*.

**Figure 6:**
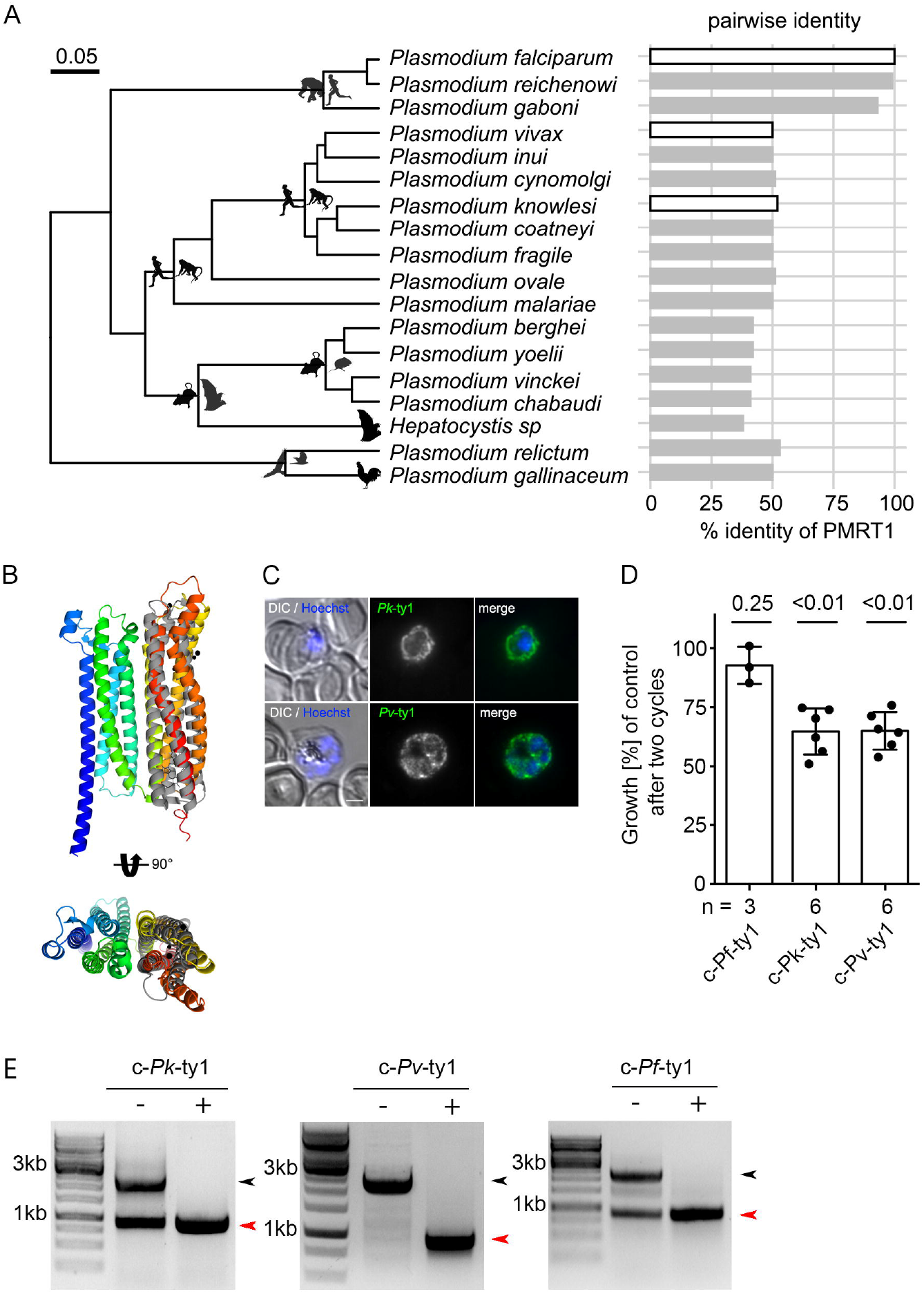
PMRT1 is a genus-specific transporter with conserved function. **(A)** Phylogenetic tree of haemosporidian parasites (modified from (95)) containing PMRT1 homologous sequences associated with data on pairwise amino acid sequence identity to *Pf*PMRT1. The phylogeny is derived from Bayesian Inference using BEAST using a fully partitioned amino acid dataset and lognormal relaxed molecular clock (95). Silhouettes depict representatives of the vertebrate hosts for each lineage and white filled bars indicate pairwise identities of PMRT1 homologs used for subsequent complementation assays. **(B)** Structural alignment of predicted *Pf*PMRT1 structure with Maquette-3 protein (PDB: 5vjt) (60). Both structures have a root mean square deviation (RMSD) over the aligned α-carbon position of 3.12 over 184 residues calculated in PyMol. **(C)** IFA of c-^*nmd3*^*Pk*-ty1 and c-^*nmd3*^*Pv*-ty1 parasites to verify correct localization of the expressed complementation fusion proteins at the parasite plasma membrane. Nuclei were stained with Hoechst-33342. Scale bar, 2 µm. **(D)** Growth of condΔPMRT1 parasites complemented with *Pf*PMRT1 homologs from *P. vivax* (PVP01_0936100) and *P. knowlesi* (PKNH_0933400). Shown are relative parasitemia values, which were obtained by dividing the parasitemia of rapalog treated cultures by the parasitemia of the corresponding untreated controls together with means +/- SD from three c-^*nmd3*^*Pf*-ty1 (≙ c-^*nmd3*^*Pf*PMRT1-ty1 Figure 3D, S5B) and six (c-^*nmd3*^*Pk*-ty1, c-^*nmd3*^*Pv*-ty1) independent growth experiments. One sample t-test **(E)** Diagnostic PCR to verify the excision of *Pf*PMRT1 on genomic level at 48 hpi for c-^*nmd3*^*Pf*-ty1, c-^*nmd3*^*Pk*-ty1 and c-^*nmd3*^*Pv*-ty1 parasites. Black arrow head, original locus; red arrow head, excised locus.

## Discussion

In this manuscript we functionally described four so called “orphan transporter” (31) in *P. falciparum*, which were partially characterized in *P. berghei*, and included two additional so far uncharacterized proteins with transporter sequence signature.

We localized *Pf*FVRT1-GFP – annotated on PlasmoDB (38) as putative divalent metal transporter – at the food vacuole of the parasite, which is in line with a previously predicted food vacuole association (1) and its reported homology (1, 61) to the conserved eukaryotic endosomal/lysosomal natural resistance-associated macrophage protein (NRAMP) transporter (62) in our structure similarity search. Repeated attempts to generate a TGD cell line failed, indicating an important role of this transporter during asexual blood stage development which is in agreement with data from a *P. falciparum* genome wide essentiality screen (63).

In concordance with recently published data identifying *Pb*DMT2 and *Pb*MFS6 as leaderless apicoplast transporters (32), we localized GFP-fusion proteins of *Pf*DMT2 and *Pf*MFS6 at the apicoplast. Successful knockdown of *Pf*DMT2 resulted in a growth defect in the second cycle after induction, resembling the described delayed death phenotype of other apicoplast genes that were functionally inactivated (32, 64–66). It suggests an essential role of *Pf*DMT2 in apicoplast physiology, as observed by Sayers *et al*. (32) for the rodent malaria *P. berghei*. This is further supported by our failed attempts to disrupt this gene using the SLI system.

We also failed to disrupt the *Pf*MFS6 locus, which is in agreement with the gene knockout studies in *P. berghei* that led to a markedly decreased multiplication rate (31, 32, 67). Nevertheless, glmS-based knock-down, although comparable to *Pf*DMT2-GFP knockdown (72.7% versus 76.8% reduction in GFP fluorescence, respectively) had no effect on parasite proliferation in our study. This might indicate that these reduced levels of *Pf*MFS6, in contrast to reduced levels of *Pf*DMT2, are sufficient for normal asexual replication *in vitro*.

Another candidate, *Pf*CDF, annotated as putative cation diffusion facilitator family protein, showed multiple cytosolic foci within the parasite with no co-localization with apical organelle markers. The homologue in *Toxoplasma gondii, Tg*ZnT (TgGT1_251630) shows a similar cellular distribution (68). It has recently been shown to transport Zn^2+^, to localize to vesicles at the plant-like vacuole in extracellular tachyzoites and to be present at dispersed vesicles throughout the cytoplasm of intracellular tachyzoites (68). The essentiality of *Pf*CDF for *in vitro* blood stage growth is debatable. We were not able to generate a clonal wild-type free TGD cell line although correct integration of the plasmid into the genomic locus could be verified (Figure S3C). This points towards its dispensability for *in vitro* blood stage growth, which is supported by i) its high (1.0) mutagenesis index score in a *P. falciparum* genome-wide mutagenesis screen (63) and ii) gene deletion experiments in rodent malaria species showing that CDF proteins are non-essential for *in vivo* blood stage development in *P. yoelii* (69) and *P. berghei* (31, 67).

Finally, two putative transporters, *Pf*ZIP1 and *Pf*PMRT1, localized to the PPM. We show that *Pf*ZIP1 is non-essential for *P. falciparum in vitro* blood stage development, in line with a high (0.7) mutagenesis index score in a *P. falciparum* genome-wide mutagenesis screen (63). However, this is in contrast to the reported strong fitness loss in *P. berghei* (67) knockout mutants and failed knockout attempts in *P. yoelli* and *P. berghei in vivo* mouse models (32, 69). These observations may reflect differences between *Plasmodium* species or differing requirements for *in vitro* and *in vivo* growth conditions.

*Pf*PMRT1 is annotated as a conserved *Plasmodium* membrane protein with unknown function. It has been described as a protein showing structural characteristics of a transporter, without sharing sequence similarities with known or putative transporters and/or conserved domains of known transporter families (2, 5). It encompasses 410 amino acids with eight predicted (70) transmembrane domains (TM) (Figure S1). The N- and C-terminal parts of *Pf*PMRT1 are both predicted (71) to be facing the cytosolic side of the parasite. Surface electrostatics indicate a clear polarity of *Pf*PMRT1 with negative charges facing the parasitophorous vacuole (PV) lumen and positive charges inside the parasite cytosol (Figure S8F). The loops protruding into the PV lumen of *Pf*PMRT1 are generally larger than the cytosolic loops and possess stretches of negatively charged amino acids likely relevant for its transport function. Further functional characterization of *Pf*MRT1 will deliver insight into its transporter capabilities and its physiological role.

Our phylogenetic analysis confirmed PMRT1 as unique for *Plasmodium* species with high sequence conservation only within the *Laverania* subgenus (72). In line with data from genome-wide mutagenesis screens (63, 67) and reported failed knockout attempts in *P. yoelii* (69), we found that *Pf*PMRT1 is essential for parasite growth, as its functional inactivation resulted in growth arrest at the trophozoite stage accompanied by the accumulation of PPM-derived protrusions within the parasite. In contrast, conditional knockdown resulted only in a growth delay, indicating that minor residual *Pf*PMRT1 protein levels appear to be sufficient to promote parasite growth. This finding was validated by episomal expression of an allelic copy under the control of the weak *sf3a2* promoter (55) in the *Pf*PMRT1 knockout parasites. Additionally, we found that *Pf*PMRT1 is essential for early gametocytogenesis. Interestingly, the induction of the knockout at stage II–III had no effect on gametocytogenesis. This might be due to sufficient amounts of *Pf*PMRT1 already present at the PPM, but could also indicate that the function of the transporter is not required for later stage gametocyte maturation.

For future work, further functional and pharmacological characterization of this transporter will provide insights into its biological role in different stages of the parasites life cycle, as transcriptomic data indicates – along with expression in blood stages (34, 35) – *Pf*PMRT1 is expressed in oocysts of *P. falciparum* (73, 74) and *P. berghei* (75).

## Material and methods

### Cloning of plasmid constructs for parasite transfection

For endogenous tagging using the SLI system (41) a 889 bp (for *Pf*PMRT1; PF3D7_1135300), 905 bp (*Pf*FVRT1; PF3D7_0523800), 827bp (*Pf*ZIP1; PF3D7_0609100), 873 bp (*Pf*DMT2; PF3D7_0716900), 877 bp (*Pf*MFS6; PF3D7_1440800), 785 bp (*Pf*CDF; PF3D7_0715900) long homology region (HR) was amplified using 3D7 gDNA and cloned into pSLI-GFP-glmS (76) (derived from pSLI-GFP (41)) using the NotI/MluI restriction site. In order to generate *Pf*PMRT1-2xFKBP-GFP a 1000 bp long HR was amplified using 3D7 gDNA and cloned into pSLI-2xFKBP-GFP (41).

For SLI-based targeted gene disruption (SLI-TGD) (41) a 501 bp (*Pf*PMRT1), 378 bp (*Pf*FVRT1), 511 bp (*Pf*ZIP1), 399 bp (*Pf*DMT2), 396 bp (*Pf*MFS6), 741 bp (*Pf*CDF) long homology region was amplified using 3D7 gDNA and cloned into the pSLI-TGD plasmid (41) using NotI and MluI restriction sites.

For conditional deletion of *Pf*PMRT1, the first 492 bp of the *Pf*PMRT1 gene were PCR amplified to append a first loxP site and a recodonized T2A skip peptide. The recodonized full-length coding region of *Pf*PMRT1 was synthesized (GenScript, Piscataway, NJ, USA) and PCR amplified with primers to add a second loxP site after the gene to obtain a second fragment. Both fragments were cloned into pSLI-3xHA (55), using NotI/SpeI and AvrII/XmaI sites. This resulted in plasmid pSLI-*Pf*PMRT1-loxP and the resulting transgenic cell line after successful genomic modification was transfected with pSkip-Flox (41) using 2 μg/ml Blasticidin S to obtain a line expressing the DiCre fragments (condΔPMRT1).

For complementation constructs, the recodonized *Pf*PMRT1 gene was PCR amplified using primers to append the TY1 sequence and cloned via XhoI and AvrII or KpnI into pEXP1comp (55) containing yDHODH as a resistance marker and different promoters (*nmd3 (*PF3D7_0729300), *sf3a2 (*PF3D7_0619900)) driving expression of the expression cassette. This resulted in plasmids c-^*nmdr*^*Pf*PMRT1-ty1 and c-^*sf3a2*^*Pf*PMRT1-ty1.

*Pf*PMRT1 homologues of *P. vivax (*PVP01_0936100) (77) *and P. knowlesi (*PKNH_0933400) (78) were amplified from parasite gDNA and cloned into p^*nmd3*^EXP1comp (55) via the XhoI/AvrII restriction site. For co-localization experiments the plasmids pLyn-FRB-mCherry (41), P40PX-mCherry (44), pARL-^*crt*^ACP-mCherry (46), pARL-^*ama1*^ARO-mCherry (49) and pARL-^*ama1*^AMA1-mCherry (51) were used. For conditional gametocyte induction yDHODH was amplified by PCR from pARL-^*ama1*^AMA1-mCherry-yDHODH (51) and cloned into GDV1-GFP-DD-hDHFR(59)(59) using the XhoI/XhoI restriction site.

Oligonucleotides and plasmids used in this study are listed in Table S1A and S1B.

### *P. falciparum* culture and transfection

Blood stages of *P. falciparum* 3D7 were cultured in human erythrocytes (O+). Cultures were maintained at 37°C in an atmosphere of 1% O_2_, 5% CO_2_ and 94% N_2_ using RPMI complete medium containing 0.5% Albumax according to standard protocols (79). To maintain synchronized parasites, cultures were treated with 5% sorbitol (80).

Induction of gametocytogenesis was done as previously described (58, 59). Briefly, GDV1-GFP-DD expression was achieved by addition of 4 µM shield-1 to the culture medium and gametocyte cultures were treated with 50 mM N-acetyl-D-glucosamine (GlcNAc) for five days starting 72 hours post shield-1 addition to eliminate asexual parasites(81). Alternatively, asexual ring stage cultures with >10% parasitemia were synchronized with Sorbitol (80) cultured for 24 hours and treated with 50 mM N-acetyl-D-glucosamine (GlcNAc) (81) for five days.

For transfection, Percoll-purified (82) late-schizont-stage parasites were transfected with 50 µg of plasmid DNA using Amaxa Nucleofector 2b (Lonza, Switzerland) as previously described(83). Transfectants were selected either using 4 nM WR99210 (Jacobus Pharmaceuticals), 2 μg/ml Blasticidin S (Life Technologies, USA), or 0.9 μM DSM1 (84) (BEI Resources; https://www.beiresources.org). In order to select for parasites carrying the genomic modification using the SLI system (41), G418 (Sigma-Aldrich, St. Louis, MO) at a final concentration of 400 µg/ml was added to 5% parasitemia culture. The selection process and testing for integration were performed as previously described (41).

For SLI-TGD, a total of six (*Pf*PMRT1, *Pf*DMT2, *Pf*ZIP1, *Pf*CDF) or eight (*Pf*FVRT1, *Pf*MFS6) independent 5 ml cultures containing the episomal plasmid were selected under G418 for at least eight weeks.

### Imaging and immunofluorescence analysis (IFA)

Fluorescence images of infected erythrocytes were observed and captured using a Zeiss Axioskop 2plus microscope with a Hamamatsu Digital camera (Model C4742-95), a Leica D6B fluorescence microscope equipped with a Leica DFC9000 GT camera and a Leica Plan Apochromat 100x/1.4 oil objective or an Olympus FV3000 with a x100 MPLAPON oil objective (NA 1.4). Confocal microscopy was performed using a Leica SP8 microscope with laser excitation at 405 nm, 490 nm, and 550 nm for DAPI, GFP, and mCherry excitation, respectively. An HC PL APO 63x NA 1.4 oil immersion objective was used and images were acquired with the HyVolution mode of the LASX microscopy software. After recording, images were deconvolved using Huygens (express deconvolution, setting ‘Standard’).

Microscopy of unfixed IEs was performed as previously described (85). Briefly, parasites were incubated in RPMI1640 culture medium with Hoechst-33342 (Invitrogen) for 15 minutes at 37°C prior to imaging. 7 µl of IEs were added on a glass slide and covered with a cover slip. Control images of 3D7 wild type parasites across the IDC are included in Figure S8D, E. BODIPY TR C5 ceramide (Invitrogen) staining was performed by adding the dye to 32 hours post invasion parasites in a final concentration of 2.5 μM in RPMI as previously described (85). For DHE staining of the parasite cytosol (44), 80 µl of resuspended parasite culture were incubated with DHE at a final concentration of 4.5 µg/ml in the dark for 15 minutes prior to imaging.

IFAs were performed as described previously (86). Briefly, IEs were smeared on slides and air-dried. Cells were fixed in 100% ice cold methanol for 3 minutes at -20°C. Afterwards, cells were blocked with 5% milk powder for 30 minutes. Next primary antibodies were diluted in PBS/3% milk powder and incubated for 2 hours, followed by three washing steps in PBS. Secondary antibodies were applied for 2 hours in PBS/3% milk powder containing 1 μg/ml Hoechst-33342 (Invitrogen) or DAPI (Roche) for nuclei staining, followed by 3 washes with PBS. One drop of mounting medium (Mowiol 4-88 (Calbiochem)) was added and the slide sealed with a coverslip for imaging.

To assess the localisation of the endogenously HA-tagged *Pf*PMRT1 IFAs were performed in suspension with Compound 2-stalled schizonts (87) to distinguish protein located at the PPM from that located at the PVM as previously done (55, 88). For this, trophozoite stages were treated with Compound 2 (1 μM) overnight, and arrested schizonts were harvested, washed in PBS, and fixed with 4% paraformaldehyde/0.0075% glutaraldehyde in PBS. Cells were permeabilized with 0.5% Triton X-100 in PBS, blocked with 3% BSA in PBS, and incubated overnight with primary antibodies diluted in 3% BSA in PBS. Cells were washed 3 times with PBS and incubated for 1 hour with Alexa 488 nm or Alexa 594 nm conjugated secondary antibodies specific for human and rat IgG (Invitrogen) diluted 1:2,000 in 3% BSA in PBS and containing 1 μg/ml DAPI. Cells were directly imaged after washing 5 times with PBS

Antisera used: 1:200 mouse anti-GFP clones 7.1 and 13.1 (Roche), 1:500 rat anti-HA clone 3F10 (Roche), 1:1000 human anti-MSP1 (89), 1:10000 mouse anti-TY1 (ThermoFischer Scientific Cat.No: MA5-23513). Contrast and intensities were linear adjusted if necessary and cropped images were assembled as panels using Fiji (90), Corel Photo-Paint X6 and Adobe Photoshop CC 2021.

## Immunoblots

For immunoblotting parasites were released from erythrocytes by incubation with 0.03% saponin in PBS for 10 minutes on ice followed by three wash steps with D-PBS. Proteins were then extracted with lysis buffer (4 % SDS, 0.5 % Triton X-100, 0.5x D-PBS in dH_2_O) in the presence of protease cocktail inhibitor (Roche) and 1 mM PMSF followed by addition of reducing SDS sample buffer and 5 minutes incubation at 55°C. Parasite proteins were separated on a 10% SDS-PAGE gel using standard procedures and transferred to a nitrocellulose membrane (Amersham(tm)Protran(tm) 0.45 µm NC, GE Healthcare) using a transblot device (Bio-Rad) according to manufacturer’s instructions or to a nitrocellulose membrane (Licor) in a tankblot device (Bio-Rad) using transfer buffer (0.192 M glycine, 0.1% SDS, 25 mM Tris-HCl pH = 8.0) with 20% methanol.

Rabbit anti-aldolase (91) and anti-SBP1 (91) antibodies were diluted 1:2,000, mouse anti-GFP clones 7.1 and 13.1 (Roche) antibody was diluted 1:500 or 1:1,000, mouse anti-Ty1 (Sigma) was diluted 1:20000, rabbit anti-BIP (92) was diluted 1:2500 and rat anti-HA clone 3F10 (Roche) antibody was diluted 1:1,000.

The chemiluminescent signal of the HRP-coupled secondary antibodies (Dianova) was visualized using a Chemi Doc XRS imaging system (Bio-Rad) and processed with Image Lab Software 5.2 (Bio-Rad). To perform loading controls and ensure equal loading of parasite material anti-aldolase antibodies were used. The corresponding immunoblots were incubated two times in stripping buffer (0.2 M glycine, 50 mM DTT, 0.05% Tween 20) at 55°C for 1 hour and washed 3 times with TBS for 10 minutes. For Western blots shown in Figure S8C fluorescent signals of secondary goat anti-rabbit IgG coupled to IRDye® 680CW and goat anti-mouse IgG coupled to IRDye® 800CW were visualized using Odyssey Fc Imager by LI-COR Biosciences.

### Growth Assay

A flow cytometry-based assay adapted from previously published assays (44, 93) was performed. For this, parasite cultures were resuspended and 20 µl samples were transferred to an Eppendorf tube. 80 µl RPMI containing Hoechst-33342 and dihydroethidium (DHE) was added to obtain final concentrations of 5 µg/ml and 4.5 µg/ml, respectively. Samples were incubated for 20 minutes (protected from UV light) at room temperature, and parasitemia was determined using an LSRII flow cytometer by counting 100,000 events using the FACSDiva software (BD Biosciences) or using an ACEA NovoCyte flow cytometer.

### Stage distribution assay

In order to obtain tightly synchronized parasite cultures, percoll purified schizonts (82) were cultured for four hours together with fresh erythrocytes, followed by sorbitol synchronization and resulting in a four-hour age window of parasites. Next, the culture was divided in four dishes and rapalog was added at a final concentration of 250 nM immediately to one dish and at 20 hours post invasion (hpi) and 32 hpi to the respective dishes. Giemsa smears and samples for flow cytometry were collected at the indicated timepoints. The parasitemia was determined using a flow cytometry assay and the stages were determined microscopically counting at least 50 infected erythrocytes per sample and timepoint.

### Gametocyte stage distribution assay

Giemsa-stained blood smears 10 days post induction of GDV1 expression were obtained and at least 10 fields of view were recorded using a 63x objective per treatment and time point. Erythrocyte numbers were then determined using the automated Parasitemia software (http://www.gburri.org/parasitemia/) while the number of gametocytes, pycnotic and asexual parasites was determined manually in >1800 erythrocytes per sample. This assay was done blinded.

### GlmS-based knockdown

GlmS based knockdown assay was adapted from previously published assays (42, 76). To induce knockdown 2.5 or 5 mM glucosamine was added to highly synchronous early rings stage parasites. As a control, the same amount of glucosamine was also added to 3D7 wildtype parasites. For all analyses, the growth medium was changed daily, and fresh glucosamine were added every day.

Knockdown was quantified by fluorescence live cell microscopy at day 1 and 3 of the growth assay. Parasites with similar size were imaged, and fluorescence was captured with the same acquisition settings to obtain comparable measurements of the fluorescence intensity. Fluorescence intensity (integrated density) was measured with Fiji(90), and background was subtracted in each image. The data were analyzed with Graph Pad Prism version 8.

GlmS based knockdown experiments in gametocytes were performed as described previously (94). Briefly, synchronized ring stage cultures were induced by the addition of shield-1. At day 3 post induction the culture was spilt into two dishes and one dish was cultured in the presence of 2.5 mM glucosamine for the remaining ten days. Knockdown was quantified by fluorescence live cell microscopy at day 7 and 10 post induction, as described above and gametocyte parasitemia was determined at day 10 post induction using the automated Parasitemia software (http://www.gburri.org/parasitemia/).

### DiCre mediated conditional knockout

The parasites containing the integrated pSLI-*Pf*PMRT1-loxP construct were transfected with pSkip-Flox (41) using 2 μg/ml Blasticidin S to obtain a line expressing the DiCre fragments. To induce excision, the tightly synchronized parasites (detailed description see growth assay) were split into 2 dishes and rapalog was added to one dish (Clontech, Mountain View, CA) to a final concentration of 250 nM. The untreated dish served as control culture. Excision was verified at genomic level after 24 and 48 hours of cultivation by PCR and on protein level by Western blot using anti-HA antibodies.

### Phylogenetic analysis

A blastp search of the PMRT1 sequence (PlasmoDB (38): PF3D7_1135300; UniProt: Q8II12) was performed against the nr database (9 May 2021) using Geneious Prime 2021.2.2 (https://www.geneious.com) and an E-value of 10e-0 (BLOSUM62 substitution matrix). Blast hits were filtered for sequences from taxa represented in the currently favored haemosporidian parasite phylogeny (95). The phylogeny derived from an amino acid alignment using Bayesian framework with a partitioned supermatrix and a relaxed molecular clock (18_amino_acid_partitioned_BEAST_relaxed_clock_no_outgroup.tre; (95)) was visualized with associated data using the R package ggtree v3.3.0.900 (96, 97). A multiple protein sequence alignment of PMRT1 and homologous sequences was performed using MAFFT v7.490 (98) using the G-INS-I algorithm to obtain a highly accurate alignment. Protein statistics were calculated using Geneious Prime 2021.2.2 (https://www.geneious.com) and EMBOSS pepstats v6.6.0.0 (99).

### Prediction of protein structures

AlphaFold structure predictions (39) were retrieved from https://alphafold.ebi.ac.uk and the PDB used for DALI protein structure homology search (40). PyMOL Molecular Graphics System, Version 2.5.2 Schrödinger was used for visualization of all structures, generation of figures and the calculation of the root mean square deviation (RMSD) between the predicted crystal structure of *Pf*PMRT1 and the Maquette-3 protein (PDB: 5vjt (60)) by cealign. The Adaptive Poisson-Boltzmann Solver (APBS) within PyMOL was used to predict the surface electrostatics of *Pf*PMRT1.

Parasite icons were generated using BioRender (biorender.com), plasmids and oligonucleotides were designed using ApE (100) and statistical analysis was performed using GraphPad Prism version 8 (GraphPad Software, USA).

## Acknowledgements

We thank Michael Filarsky for providing the pHcamGDV1-GFP-DD_hDHFR plasmid, Egbert Tannich for providing *P. vivax* and *P. knowlesi* gDNA, Mike Blackman for the anti-MSP1 antibody, Jacobus Pharmaceuticals for WR99210, Greg Burri for the parasitemia software and the Advanced Light and Fluorescence Microscopy (ALFM) facility at the Centre for Structural Systems Biology (CSSB), in particular Roland Thuenauer, for support with light microscopy image recording and analysis. DSM1 (MRA-1161) was obtained from MR4/BEI Resources, NIAID, NIH. Furthermore, we thank Maria Rosenthal for her help with the visualization of the predicted protein structures.

## Author contribution

Conceptualization: JSW, TWG, AB, JSTR

Methodology: GN, SG

Investigation: JSW, PMR, JY, GF; JSTÄ, HVT, AA, IH, BL

Formal Analysis: JSTR

Writing original manuscript: JSW, AB, JSTR, DW

Review & Editing: JSW, PMR, TWG, AB, JSTR, DW

Funding Acquisition: DW, CL, TWG, AB, JSTR

Resources: TWG

Project Administration: TWG, AB, JSTR

Supervision: DW, TS, SG, TWG, AB

All authors read and approved the manuscript.

## Figures

**Figure S1: Structure predictions and structure homology search of candidate proteins**

**(A)** AlphaFold structure predictions of the six selected orphan transporters visualized in PyMol. **(B)** Results from protein structure comparison server Dali using the AlphaFold-generated PDB files of the selected transporters as input structure. Shown are the top five non-redundant hits with Z score (significance estimate), msd (difference between the root-mean-square-deviation (rmsd) value associated with a protein structure pair and the rmsd value that would have been observed in the case that the two structures had the same crystallographic resolution), lali (number of aligned positions), nres (number of residues in the matched structure) and %id (the percentage sequence identity in the match).

**Figure S2: Validation of generated transgenic cell lines by PCR and Western blot**.

**(A)** Confirmatory PCR of unmodified wildtype (WT) and transgenic knock-in (KI) cell lines (PF3D7_0523800-GFP-glmS (*Pf*FVRT1), PF3D7_0609100-GFP-glmS (*Pf*ZIP1), PF3D7_0715900-GFP-glmS (*Pf*CDF), PF3D7_0716900-GFP-glmS (*Pf*DMT2), PF3D7_1440800-GFP-glmS (*Pf*MFS6) and PF3D7_1135300-GFP-glmS (*Pf*PMRT1)) to check for genomic integration at the 3’- and 5’-end of the locus. Position of the primer used are indicated with numbered arrows in Figure 1A. **(B)** Western Blot analysis of wildtype (3D7) and knock-in (KI) cell lines using mouse anti-GFP to detect the tagged full-length protein (upper panel) and rabbit anti-aldolase to control for equal loading (lower panel). Protein size is indicated in kDa. Expected molecular weight for GFP fusion proteins: *Pf*FVRT1 (107.5 kDa), *Pf*ZIP1 (69.0 kDa), *Pf*DMT2 (66.4 kDa), *Pf*MFS6 (98.8 kDa), *Pf*PMRT1 (77.5 kDa), *Pf*CDF (91.6 kDa) **(C)** Localization of *Pf*CDF-3xHA by IFA in ring, trophozoite and schizont parasites. Nuclei were stained with Hoechst. Diagnostic PCR of unmodified wildtype (WT) and transgenic knock-in (KI) cell line. **(D)** Localization of *Pf*PMRT1_2xFKBP-GFP across the IDC. Nuclei were stained with DAPI. Scale bar, 2 µm. Diagnostic PCR of unmodified wildtype (WT) and transgenic knock-in (KI) cell line. **(E)** Localization of *Pf*ZIP1-GFP in merozoites. Nuclei were stained with DAPI. Scale bar, 2 µm. **(F)** Confocal microscopy of *Pf*PMRT1-GFP co-expressing the PPM marker Lyn-mCherry. Scale bar, 1µm. Nuclei were stained with Hoechst.

**Figure S3: Targeted gene disruption (TGD) of *Pf*ZIP1 and *Pf*CDF**.

**A)** Schematic representation of TGD strategy using the selection-linked integration system (SLI). pink, human dihydrofolate dehydrogenase (hDHFR); grey, homology region (HR); green, green fluorescence protein (GFP) tag; dark grey, T2A skip peptide; blue, neomycin resistance cassette. Stars indicate stop codons, and arrows depict primers (P1 to P4) used for the integration check PCR. **(B)** Localization of *Pf*ZIP1-TGD-GFP in ring, trophozoite and schizont parasites. Nuclei were stained with Hoechst-33342. Scale bar, 2 µm. Confirmatory PCR of unmodified wildtype (WT) and transgenic targeted gene disruption (TGD) cell line. Growth curves of PfZIP1-TGD vs. 3D7 parasites monitored over five days by FACS. Three independent growth experiments were performed and a summary is shown as percentage of growth compared to 3D7 parasites. **(C)** Localization of *Pf*CDF-TGD in ring, trophozoite and schizont parasites. Nuclei were stained with DAPI. Confirmatory PCR of unmodified wildtype (WT) and transgenic targeted gene disruption (TGD) cell line. Scale bar, 1 µm.

**Figure S4: Conditional knockdown via glmS system**.

Live cell microscopy of **(A)** *Pf*FVRT1-GFP-glmS, **(B)** *Pf*CDF-GFP-glmS, **(C)** *Pf*ZIP1-GFP-glmS **(D)** *Pf*DMT2-GFP-glmS, **(E)** *Pf*MFS6-GFP-glmS and **(F)** *Pf*PMRT1-GFP-glmS parasites 40 hours after treatment without (control) or with 2.5 mM Glucosamine. Nuclei were stained with Hoechst-33342. Scale bar, 2 µm. **(G)** Individual growth curves of the growth assays shown in Figure 2D. **(H)** PCR using a GFP forward and glmS reverse primer confirming the presence of the GFP and glmS sequence in the pSLI-*Pf*FVRT1-GFP-glmS and *Pf*CDF-GFP-glmS plasmids. pSLI-PF3D7_0631900-GFP (35) was used as negative control.

**Figure S5: Conditional knockout of *Pf*PMRT1 via DiCre-based system**

Replicates of growth curves of condΔPMRT1, c-^*nmd3*^*Pf*PMRT1-ty1 and c-^*sf3a2*^*Pf*PMRT1-ty1 parasites +/- rapalog monitored over five days by FACS shown in Figure 3.

**Figure S6: Conditional knockout of *Pf*PMRT1**

**(A)** Parasite stage distribution in Giemsa smears displayed as heatmap showing percentage of stages for control, 4 hpi, 20 hpi or 32 hpi rapalog treated 4 hour window synchronized condΔPMRT1parasite cultures over one cycle. **(B)** Giemsa smears of control and 4 hpi, 20 hpi or 32 hpi rapalog treated parasites at 4, 16, 20, 24, 32, 40 and 48 hpi. Scale bar, 5 µm.

**Figure S7: Conditional knockdown of *Pf*PMRT1 has no effect during gametocyte development**.

**(A)** Confirmatory PCR of unmodified wildtype (WT) and transgenic 3D7-iGP-*Pf*PMRT1-GFP-glmS to check for genomic integration at the 3’- and 5’-end of the locus. Position of the primer used are indicated with numbered arrows in Figure 1A. **(B)** Schematic representation of the experimental setup. **(C)** Live cell microscopy of 3D7-iGP*-Pf*PMRT1-GFP stage I – V gametocytes. Scale bar, 2 µm. **(D)** Giemsa smears of stage I – V gametocytes cultured either without (control) or with 2.5 mM glucosamine. Scale bar, 5 µm. **(E)** Quantification of knockdown by measuring mean fluorescence intensity (MFI) density and size (area) of parasites at day 7 and day 12 post induction of gametocytogenesis cultured either without (control) or with 2.5 mM glucosamine. Scale bar, 2 µm. Statistics are displayed as mean +/- SD of four independent experiments and individual data points are displayed as scatterplot color-coded by experiments according to Superplots guidelines(101)(101). P-values displayed were determined with two-tailed unpaired t-test. **(F)** For each condition gametocytemia at day 10 post gametocyte induction was determined by counting between 1256-2653 (mean 2147) cells per condition in Giemsa-stained thin blood smears. Displayed are means +/- SD of independent growth experiments with the number of experiments (n) indicated. P-values displayed were determined with two-tailed unpaired t-test.

**Figure S8:** Individual growth curves of c-^*nmd3*^*Pk*-ty1 **(A)** and c-^*nmd3*^*Pv*-ty1 **(B)** parasites +/- rapalog monitored over two IDCs by FACS shown in Figure 6. **(C)** Western Blot analysis of c-^*nmd3*^Pf-ty1, c-^*nmd3*^*Pk*-ty1 and c-^*nmd3*^*Pv*-ty1 cell lines using mouse anti-ty1 to detect the tagged full-length protein (upper panel) and rabbit anti-BIP to control for loading (lower panel). Protein size is indicated in kDa. **(D)** and **(E)** 3D7 wild type parasites imaged across the IDC to establish autofluorescence levels with Zeiss Axioskop 2plus microscope (D) or Leica D6B fluorescence microscope (E). **(F)** Surface electrostatics of the predicted *Pf*PMRT1 structure generated by APBS within PyMol.

**Table S1: Oligonucleotides and plasmids used in this study**

