## Supplementary material for "PMRT1, a *Plasmodium* specific parasite plasma membrane transporter is essential for asexual and sexual blood stage development": Table S1

**Table S1:** Oligonucleotides **(A)** and plasmids **(B)** used in this study.

**A**

| internal # | Target or Primer name | Sequence | Purpose |
|---|---|---|---|
| 284 | PF3D7_0716900 rv | GGGacgcgtTAACATCAATTTTGCTTTTTTGGG | cloning GFP-glmS |
| 285 | PF3D7_0716900 fw | GGGgcggccgctaaGGGTGTGGTAATTCAAGTACG | |
| 280 | PF3D7_0523800 rv | GGGacgcgtATTTCGTTGAATATAATTTTTTTTAATTG | |
| 281 | PF3D7_0523800 fw | GGGgcggccgctaaGTTTTCTTGTTGGTATTTTAGCTG | |
| 282 | PF3D7_0609100 rv | GGGacgcgtATGATTATGACCATGATCATGATC | |
| 283 | PF3D7_0609100 fw | GGGgcggccgctaaGGATAGCAGGTGTTACGGTTTCTTTATC | |
| 288 | PF3D7_1440800 rv | GGGacgcgtATTAGTAATAGAATTTTTCATCTTG | |
| 289 | PF3D7_1440800 fw | GGGgcggccgctaaCATTTGCTTCAAATTTGATGAG | |
| 290 | Pf3D7_1135300 rv | GGGacgcgtAGAAGTTTTTGGGGCATATTTCTTTG | |
| 338 | Pf3D7_0715900 (11) | GGGgcggccgctaaCTGGAAATAAAATAGATGGAACGTCTTG | |
| 339 | Pf3D7_0715900 (11) | GGGacgcgtAGTATCCCCTTTCAATGTGGAAC | |
| 291 | Pf3D7_1135300 fw | GGGgcggccgctaaGTTATATTATAAAAGGATGATTGG | |
| 315 | PF3D7_0523800 (1) TGD fw | GCGGCCGCTAAAGGAGCACTAAAGGCCAAAGGAAGT | cloning TGD |
| 316 | PF3D7_0523800 (1) TGD rv | acgcgtAATAATGTCATCCTTTTCATTATTAATATT | |
| 317 | PF3D7_0609100 (2) TGD fw | GCGGCCGCTAAGATTTGTTATTTGCAAAAATAATTTGTATT | |
| 318 | PF3D7_0609100 (2) TGD rv | acgcgtATGTctataaaaaaaataaatcacatac | |
| 319 | PF3D7_0716900 (3) TGD fw | GCGGCCGCTAAAAAAATGAAATTATATTTTGTACGACATTC | |
| 320 | PF3D7_0716900 (3) TGD rv | acgcgtGGCAATACCAAAGGTAATTAATAAAATTCC | |
| 323 | PF3D7_1440800 (5) TGD fw | GCGGCCGCTAAACAATGTTTAAGGATGATGAAAATAATTTT | |
| 324 | PF3D7_1440800 (5) TGD rv | acgcgtTGTATTAACAACAACTATAAATACAGAACT | |
| | Pf3D7_0715900 SLI-Tdg F | GGTGCGGCCGCGTGGATAAGATGTCGCGTTTG | |
| | Pf3D7_0715900 SLI-Tgd R | GGTACGCGTGTGTCCTACATTTTTATCATCTTCCTC | |
| | PF3D7_1135300 – fw NotI TGD | CTCGgcggccgctaaAAAAGTATGATCTCTGGAATATCCAG | |
| | PF3D7_1135300 – rv MluI TGD | TCCTacgcgtATTTATAAAGAGATCTGTTTTATTATC | |
| | PF3D7_1135300 fw NotI loxP | CTCGGCGGCCGCCCGGTAAATCTCTGGAATATCCAGCAAAATTG | cloning 1135300 loxp |
| | PF3D7_1135300 rv SpeI loxP: | TCCTACTAGTATTTATAAAGAGATCTGTTTTATTATC | |
| | PF3D7_1135300 fw AvrII loxP: | CTCGCCTAGGATGAAGTCAATGATAAGCGGTATTAG | |
| | PF3D7_1135300 rv XmaI loxP: | TCCTCCCGGGGCTTGTCTTAGGTGCGTACTTTTTAC | |
| 294 | PF3D7_0609100 int_fw | GGTAGTAACAACCTTTGGTTGTTTTATTCC | integration check PCR |
| 295 | PF3D7_0609100 int_rv | tatgtggaaggttaattaaatggacaaggg | |
| 296 | PF3D7_0716900 int_fw | GAAATTATATTTTGTACGACATTCATACTA | |
| 298 | PF3D7_1440800 int_fw | GCACAGAACATTTAAGAAGCAATGATTTTA | |
| 299 | PF3D7_1440800 int_rv | caacttgactagccaaaatgttggttctgg | |
| 300 | PF3D7_1135300 int_fw | GATCTCTGGAATATCCAGCAAAATTGTTG | |
| 302 | PF3D7_0716900 int_rv | atatttatttattttgaaccgataagctag | |
| 305 | PF3D7_1135300 int_rv | gccatatatattatacatattaaatataaagac | |
| 308 | PF3D7_0523800 int_fw | GGCCAAAGGAAGTTGGCTAACGGGGTGGTAG | |
| 309 | PF3D7_0523800 int_rv | gtgttcattcatttaccttttgaatgg | |
| 369 | TGD (PF3D7_0609100 ) int check fw | atatatgttttaaagcttcaaaatg | |
| 370 | TGD (PF3D7_0609100 ) int check rv | gaggagaagcaaaaacgaaagttaac | |
| 392 | TGD (Pf3D7_0715900) int check fw | CCCACCGAAATGAACTCTTCGTTGC | |
| 393 | TGD (Pf3D7_0715900) int check rv | CTTTACCTTTAGAGGAGGAATTATTAG | |
| 394 | SLI TGD it fw Adelaide | gtggaattgtgagcggataac | |
| 327 | Pf3D7_1135300 compl | ctcgagATGAAGTCAATGATAAGCGG | cloning complementation constrcuts |
| 328 | Pf3D7_1135300 compl | cctaggGCTTGTCTTAGGTGCGTACT | |
| 553 | PKNH_0933400 fw 553 | GGGctcgagATGAAGGGAACGTACGTAG | |

| | | | |
|---|---|---|---|
| 554 | PKNH_0933400 rv 554 | GGGcctaggCACCGCCTTCGAGGCGTAC | |
| 555 | PVP01_0936100 fw 555 | GGGctcgagATGAAGGGAACGTACGTCG | |
| 556 | PVP01_0936100 rv 556 | GGGcctaggCACCGCCTTCGAGGCGTAC | |
| p37 | Pf3D7_1135300 fw 5' (TKo) | GATTTTGATATATGATTATAGGATAG | excision PCR primer |
| | Neo 40 rv | CGAATAGCCTCTCCACCCAAG | |
| | Pf3D7_1135300 fw NotI | CTCGgcggccgctaaTTATTATAAATCATATAATAAAATAAATG | cloning 1135300-2xFKBP-GFP |
| | Pf3D7_1135300 rv AvrII | TCCTcctaggAGAAGTTTTTGGGGCATATTTCTTTG | |
| 443 | Pf3D7_0715900 (11) KpnI | GggtaccAGTATCCCCTTTCAATGTGGAAC | cloning 0715900-3xHA |
| | GFP 633 fw | GCCCTTTCGAAAGATCCC | confirmatory PCR |
| | glmS rv | GATTTCTCTTTGTTCAAGGAGTCACC | |

**B**

| | |
|---|---|
| pSLI-*Pf*FVRT1-GFP-glmS | this study |
| pSLI-*Pf*ZIP1-GFP-glmS | this study |
| pSLI-*Pf*CDF-GFP-glmS | this study |
| pSLI-*Pf*DMT2-GFP-glmS | this study |
| pSLI-*Pf*MFS6-GFP-glmS | this study |
| pSLI-*Pf*PMRT1-GFP-glmS | this study |
| pSLI-*Pf*FVRT1-GFP-TGD | this study |
| pSLI-*Pf*ZIP1-GFP-TGD | this study |
| pSLI-*Pf*CDF-GFP-TGD | this study |
| pSLI-*Pf*DMT2-GFP-TGD | this study |
| pSLI-*Pf*MFS6-GFP-TGD | this study |
| pSLI-*Pf*PMRT1-GFP-TGD | this study |
| pSLI-*Pf*CDF-3xHA | this study |
| pSLI-*Pf*PMRT1-2xFKBP-GFP | this study |
| P40PX-mCherry | Jonscher et al. 2018 (44) |
| pLyn-FRB-mCherry | Birnbaum et al. 2017 (41) |
| pACP-mCherry | Birnbaum et al. 2020 (46) |
| pARL-$^{ama1}$ARO-mCherry | Cabrera et al. 2012 (49) |
| pARL-$^{ama1}$AMA1-mCherry | Wichers et al. 2021 (51) |
| pSLI-*Pf*PMRT1-loxP | this study |
| pARL-$^{nmd3}$*Pf*PMRT1-ty1 | this study |
| pARL-$^{sf3a2}$*Pf*PMRT1-ty1 | this study |
| pHcamGDV1-GFP-DD-yDHODH | this study |
| pSkip-Flox | Birnbaum et al. 2017 (41) |
| pARL- $^{nmd3}$*Pv*PMRT1-ty1 | this study |
| pARL- $^{nmd3}$*Pk*PMRT1-ty1 | this study |
