## Supplementary figures and images for "PMRT1, a *Plasmodium* specific parasite plasma membrane transporter is essential for asexual and sexual blood stage development"

### Figure S1

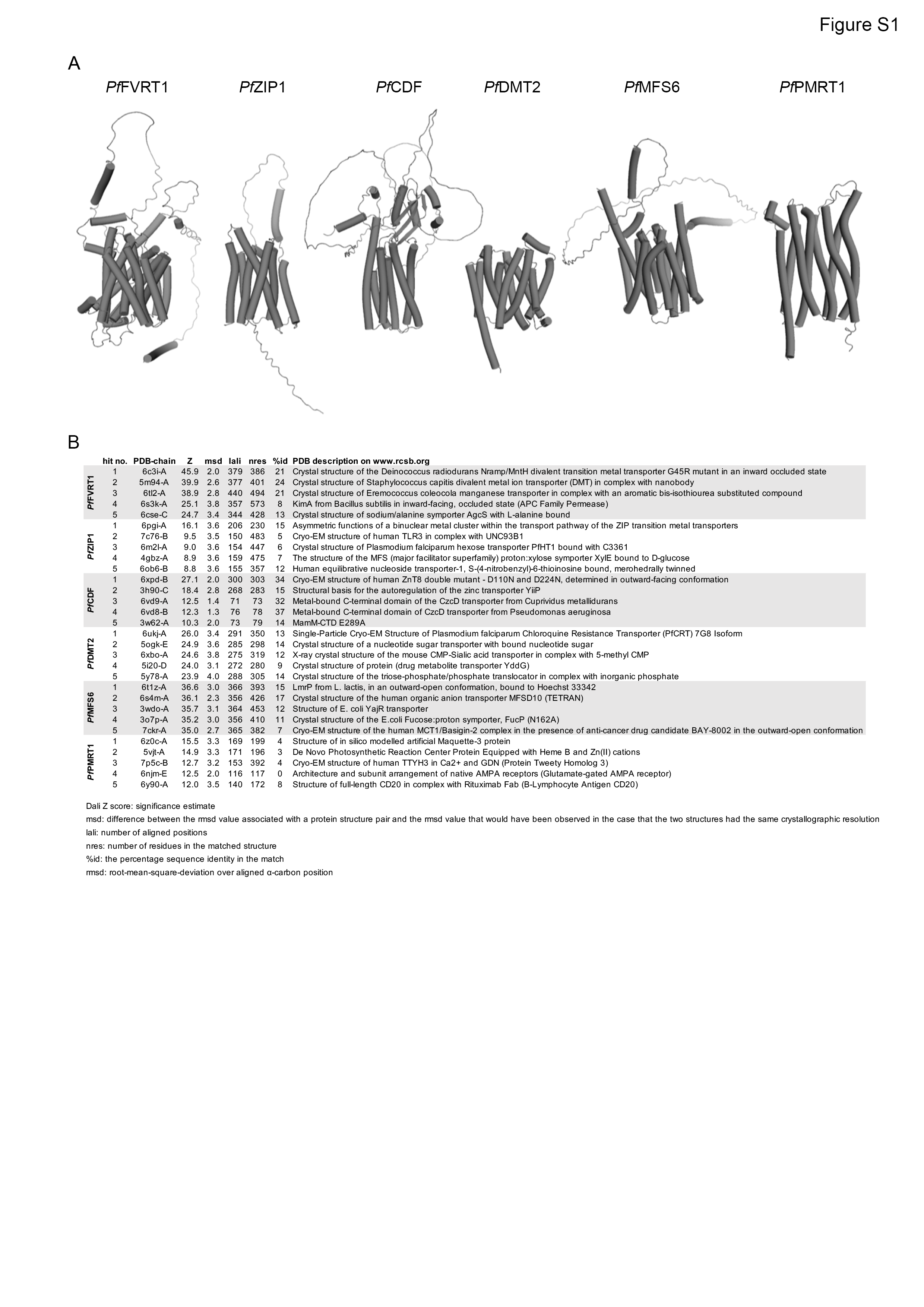

### Figure S2

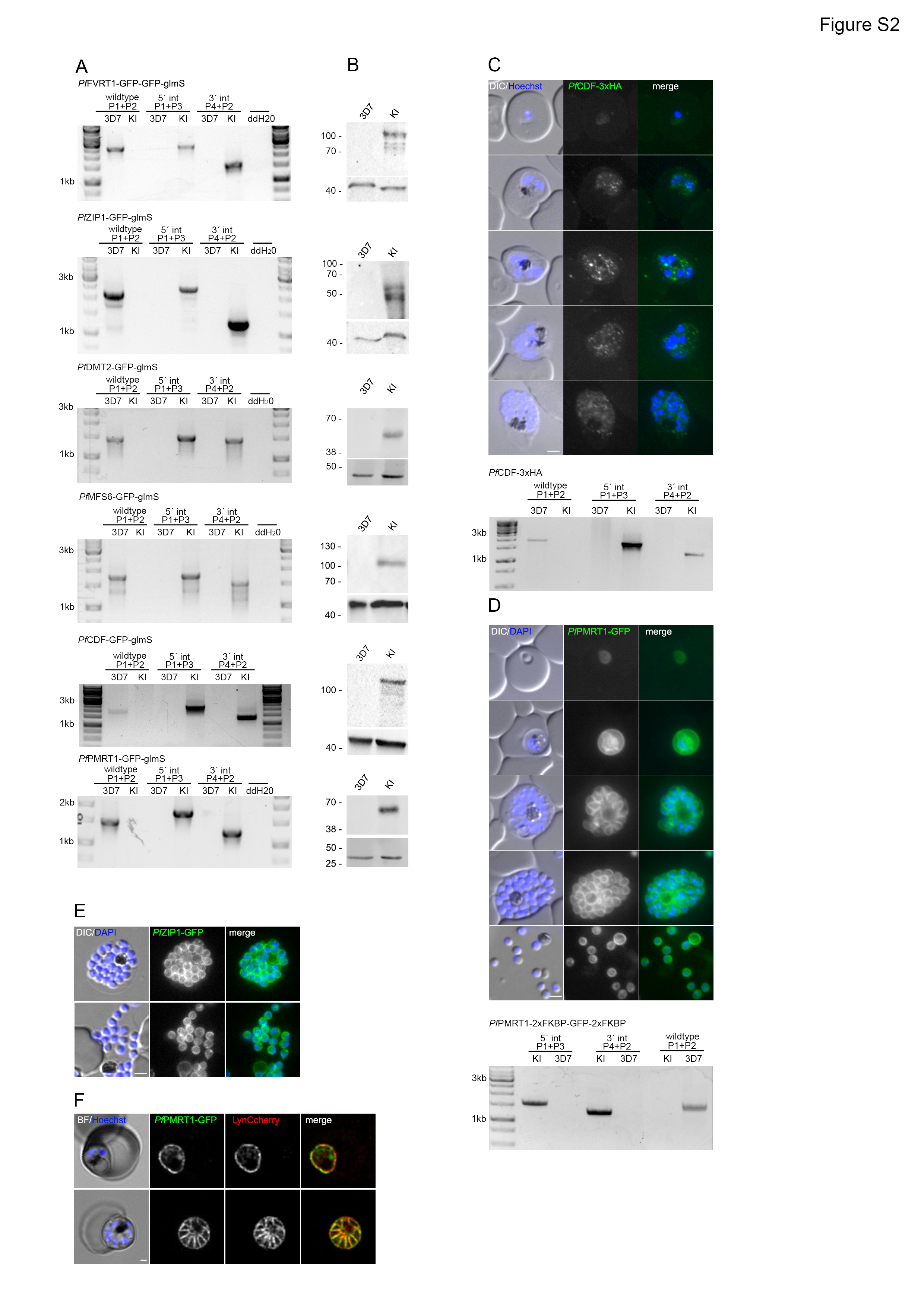

### Figure S3

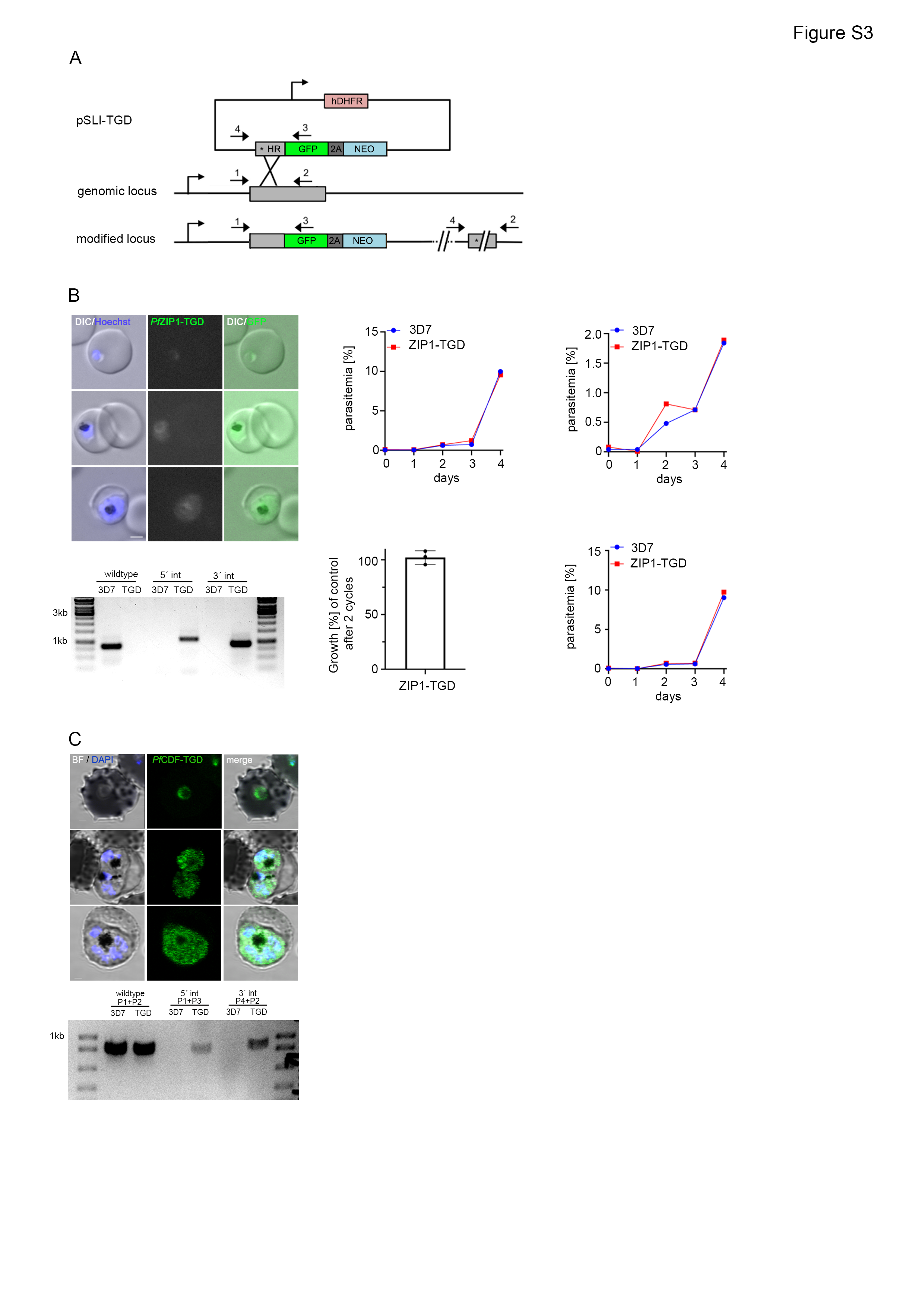

### Figure S4

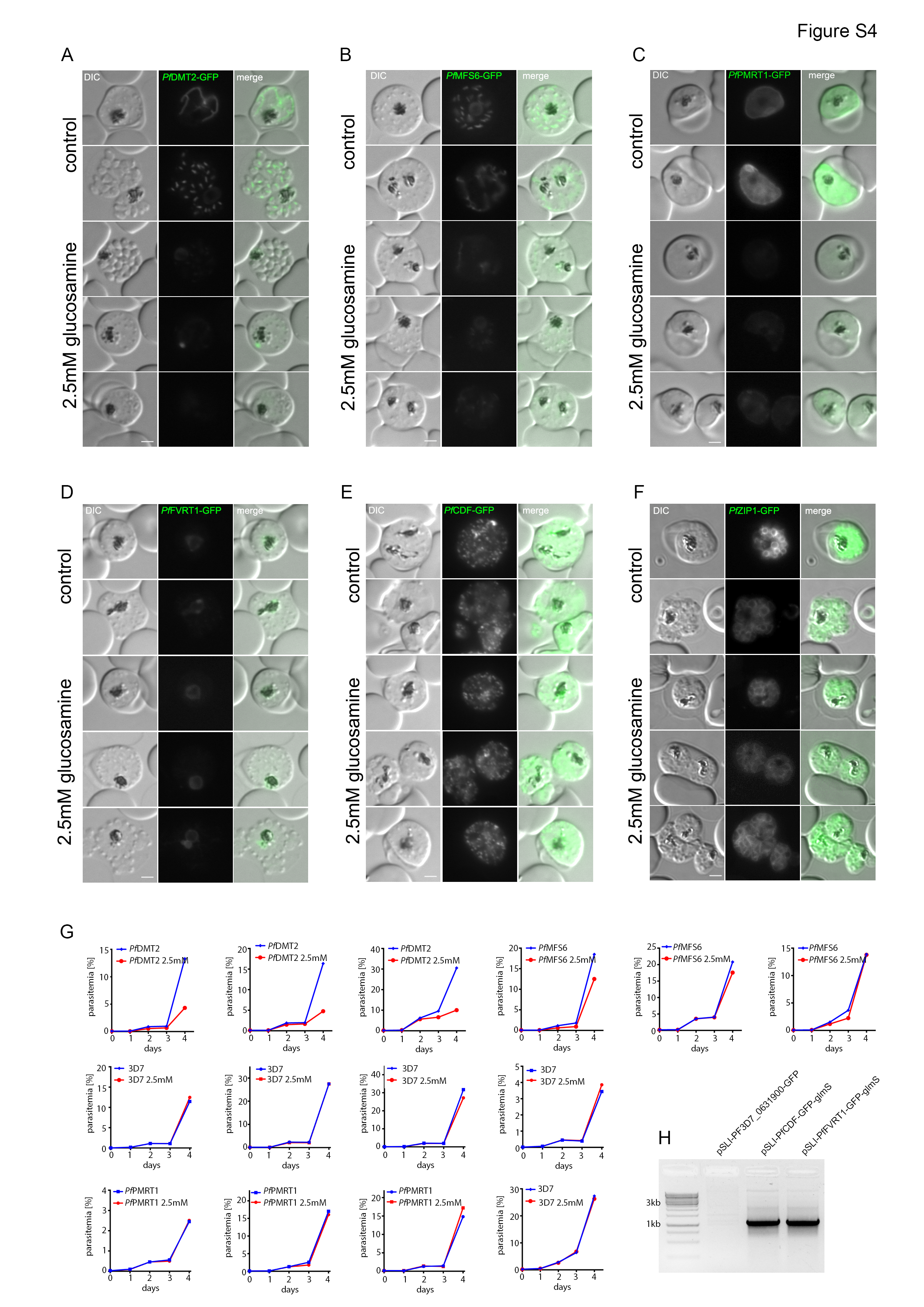

### Figure S5

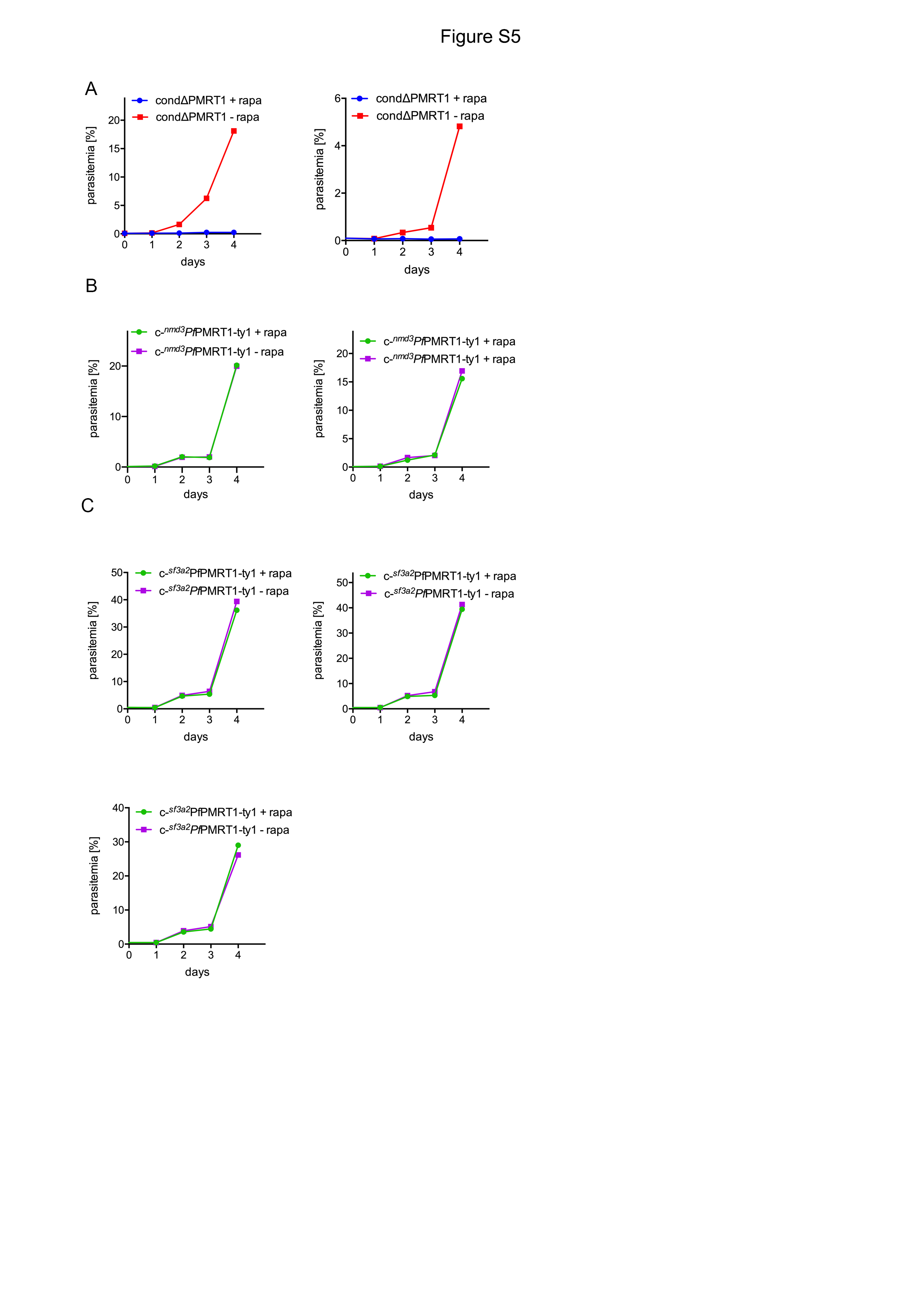

### Figure S6

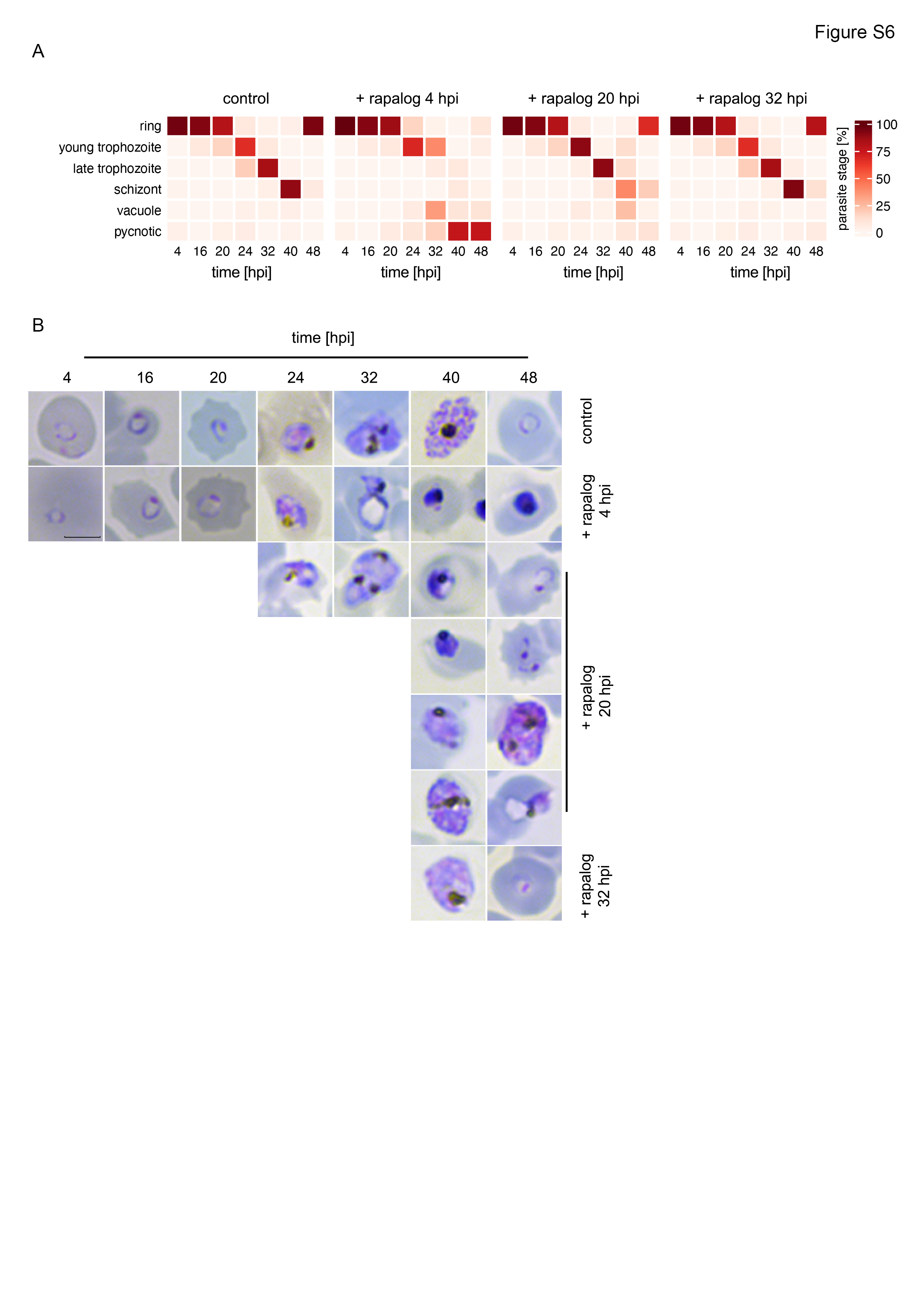

### Figure S7

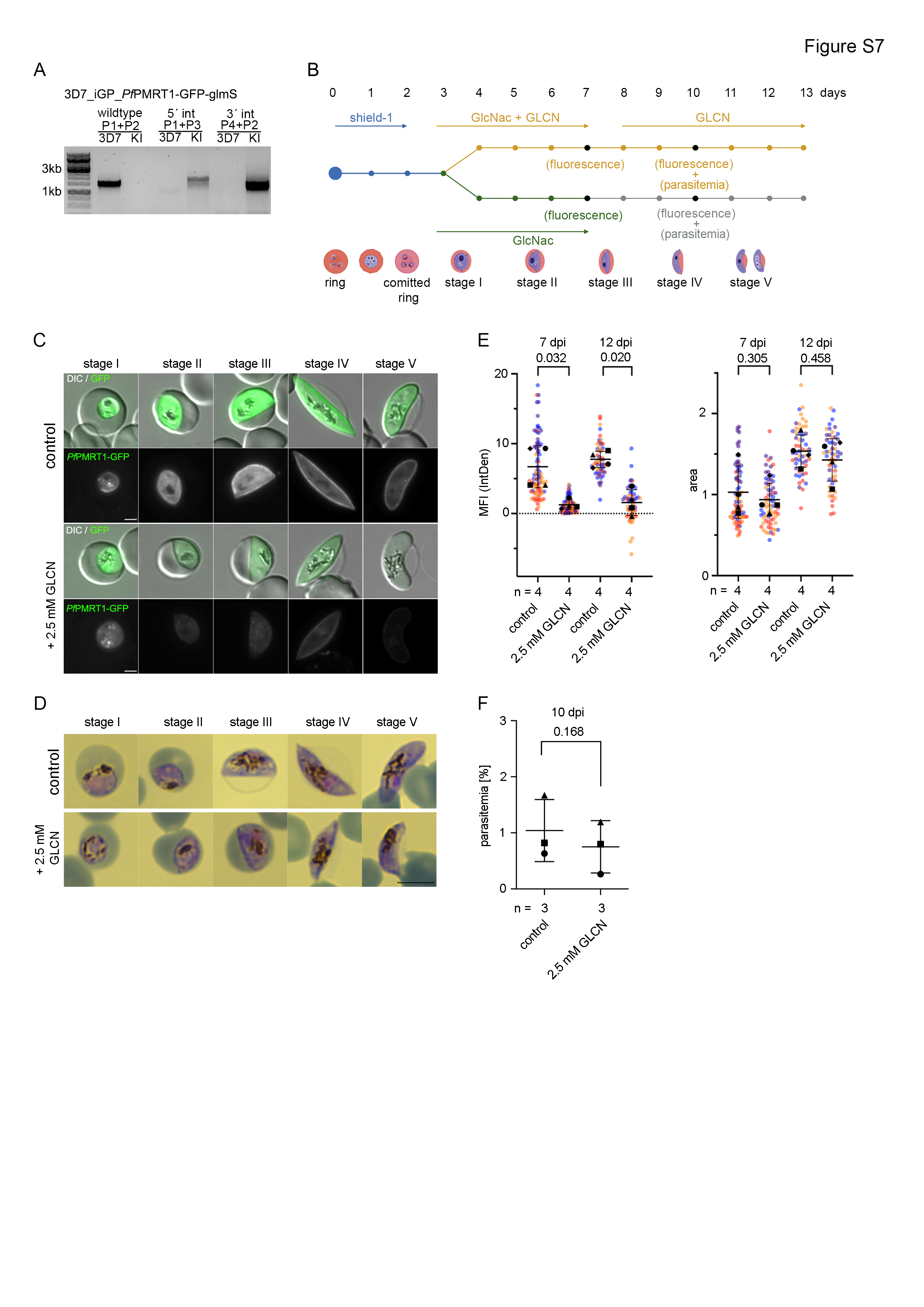

### Figure S8

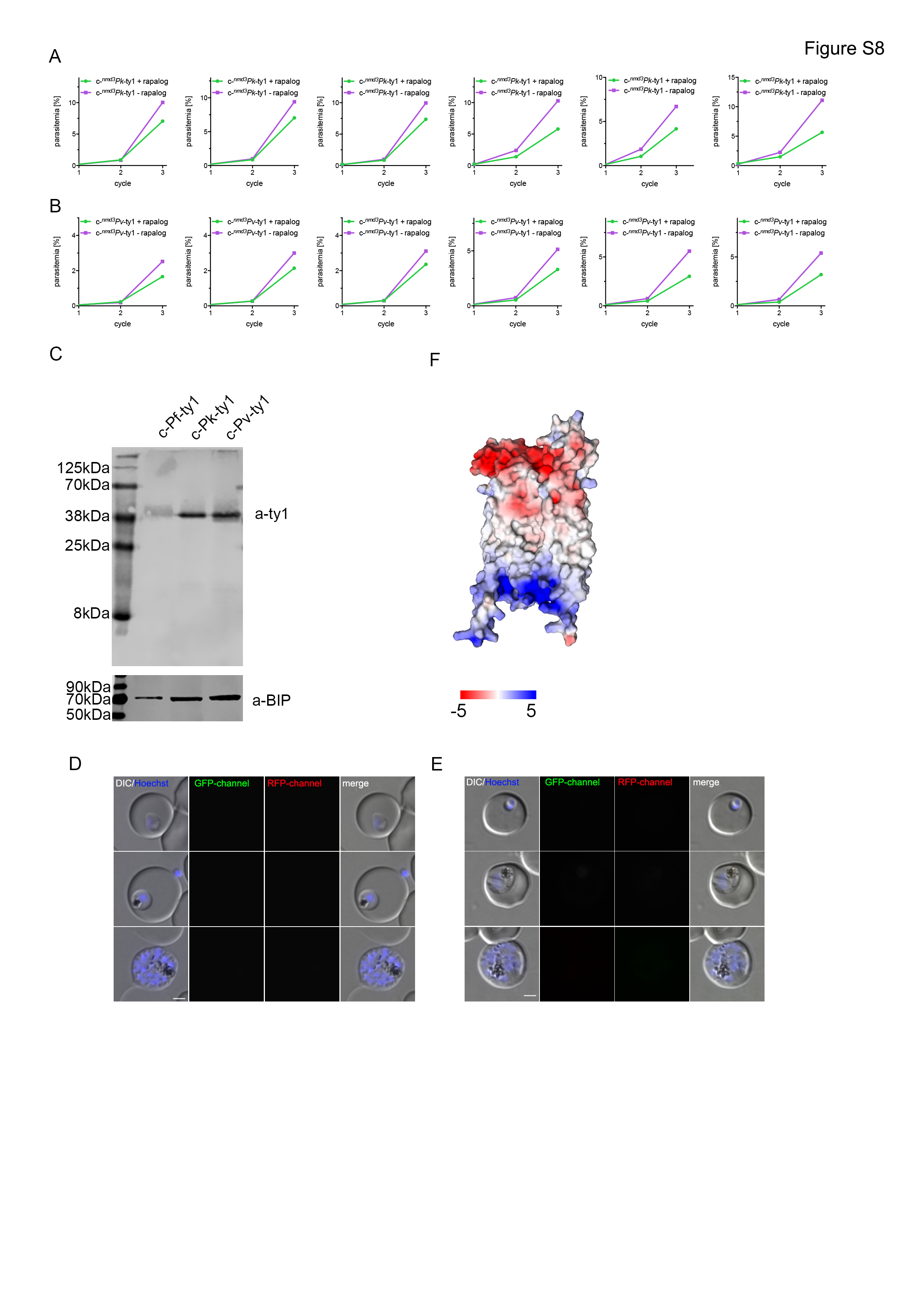
